# FoxP3-mediated blockage of ryanodine receptor 2 is the molecular basis for the contact-based suppression by regulatory T cells

**DOI:** 10.1101/2022.05.02.490213

**Authors:** Xiaobo Wang, Shuang Geng, Junchen Meng, Ning Kang, Xinyi Liu, Yanni Xu, Huiyun Lv, Ying Xu, Xun Xu, Xinrong Song, Bin Zhang, Xin Wang, Nuerdida Nuerbulati, Ze Zhang, Di Zhai, Xin Mao, Ruya Sun, Xiaoting Wang, Ruiwu Wang, Jie Guo, S. R. Wayne Chen, Xuyu Zhou, Tie Xia, Hai Qi, Xiaoyu Hu, Yan Shi

## Abstract

The suppression mechanism of regulatory T cells is an intensely investigated topic. As our focus has shifted towards a model centered on indirect inhibition of dendritic cells, a universally applicable effector mechanism controlled by FoxP3 expression has not been found. Here, we report that FoxP3 blocks the transcription of ER Ca^2+^-release channel ryanodine receptor 2. Reduced RyR2 shuts down basal Ca^2+^ oscillation in Tregs, which reduces m-Calpain activities that is needed for T cells to disengage from DCs, suggesting a persistent blockage of DC antigen presentation. RyR2 deficiency renders the CD4^+^ T cell pool to become immune suppressive, and behave in the same manner as FoxP3^+^ Tregs in viral infection, asthma, hypersensitivity, colitis and tumor development. In the absence of FoxP3, RyR2-deficient CD4^+^ T cells rescue the systemic autoimmunity associated with Scurfy mice. Therefore, FoxP3-mediated Ca^2+^ signaling inhibition may be a central effector mechanism of Treg immune suppression.

**One Sentence Summary:** Calcium channel RyR2 dictates Treg adhesion-based suppression

## INTRODUCTION

The suppressive mechanism of regulatory T cells remains a topic of continuing debate. Proposed mechanisms, including T cell cytolysis, surface protein extraction, local generation of adenosine etc. all require Treg binding to their targets of suppression (*1*). In more recent years, a string of reports has shifted our focus to the direct suppression of dendritic cells (DC). This proposal has two instinctual rationales. On the one hand, Tregs are vastly outnumbered by CD4^+^ T cells, yet retain a numerical advantage to DCs with a rough ratio of two to one *in vivo*, inferring a more logical and judicious use of suppressive power. Second, the direct DC/Treg binding appears to be commonly observed both *in vivo* and *in vitro* (*2-7*). While such binding is regarded to exert suppression on DCs, no consensus has been reached with regard to an exact mode of operation. Among proposals, Sakaguchi’s group suggests that this is mainly via the transendocytosis or trogocytosis of costimulatory molecules by CTLA-4 expressed on Tregs (*8*). Shevach’s group proposes that MHC class II extraction plays an important role in dampening DCs in antigen presentation (*9*). Our group believes that the binding between Tregs and DCs is mediated by LFA-1 and ICAM-1, and the binding itself is so exuberantly strong that DC cytoskeleton is too distorted to support antigen presentation to other T cells (*2, 7*). With time, these differences will likely be resolved to produce a more refined and comprehensive picture regarding how Tregs suppress DCs via contact. However, for the discussion to continue, one essential question needs to be answered: why do Tregs bind DCs with such a usually strong force and how this excessive force may impact DCs. Onishi et al. first used *in vitro* experiments to suggest that strong binding was mediated by LFA-1 (*4*). On that basis, we used single cell force spectroscopy (SCFS) to provide a molecular explanation how this may work. Tregs have intrinsically low m-Calpain activities. The lack of this ubiquitous calcium-regulated protease renders Tregs unable to recycle its surface LFA-1, a basic mechanism used by CD4^+^ T cells to disengage from a transient binding partner (*10-12*), Tregs are therefore locked in a perpetual state of high adhesion. DCs thus engaged use their unique actin-bundling protein Fascin-1 to polarize their cortical cytoskeleton towards Treg binding site, depriving them of their ability to form a stable contact with other T cells. As LFA-1 conformational changes and expression levels are roughly equal to other T cells, the low m-Calpain activity becomes emblematic for Tregs (*2*). Whether this feature is essential to Treg suppression and how it is regulated by the master regulation of FoxP3 are therefore interesting questions.

We report here that basal calcium oscillation in Tregs is severely depressed. mRNA expression analysis returned a list of calcium regulators with reduced abundance in Tregs. The most reduced is Ryanodine receptor 2 (RyR2), a sarcoplasmic reticulum Ca^2+^ release channel with its cytoplasmic domain in close contact to the inner leaflet of the plasma membrane (*13*). Using several model cell lines, we found that FoxP3 expression autonomously suppresses RyR2 expression, targeting a stretch of guanosine-rich sequence roughly 200 bp before RyR2 start codon. Conventional T cells with RyR2 knockdown bind to DCs with forces similar to that of Tregs and become suppressive, echoing our early findings that m-Calpain blockage imparts Treg-like inhibition to Tconvs. In addition, T cells with CD4^+^-specific RyR2 deletion become immune suppressive both *in vitro* and *in vivo* in the absence of FoxP3 expression. In the absence of Tregs, RyR2^-/-^ Tconvs restore the immune homeostasis in all aspects of immune deficiency in Scurfy mice, reminiscent of the original Sakaguchi’s reports that co-transfer of untreated CD4^+^ T cells blocked CD5^low^CD4^+^ T cells’ ability to induce systemic autoimmunity in athymic mice (*14*) and that this suppressive capacity was associated with a subpopulation of CD25^+^ T cells (*15*). Our work therefore reveals a distinct, FoxP3-mediated effector function of Tregs and may provide the molecular base of the previously elusive contact-dependent suppression by Tregs.

## RESULTS

### Reduced RyR2 activity is the basis of contact-dependent suppression by Tregs

As m-Calpain activities are reduced in Tregs, we analyzed whether the expression level is lower. Figure 1A shows that no difference in m-Calpain was found between Tconvs and Tregs at the protein or mRNA level. Calpains are regulated by distinct intracellular Ca^2+^ availability. There is a persistent reduction of overall Ca^2+^ signal in Tregs (fig. S1A). To study this in detail, we observed Ca^2+^ signals in both Tconvs and Tregs in resting state with two different indicators, Fluo-4 AM and ratiometric CAL RED R525/650 AM (*16*). While spontaneous Ca^2+^ oscillations are a common feature shared by many immune cells, including Tconvs, this activity was essentially absent in Tregs (Fig. 1, B and C), supporting their low m-Calpain activation.

**Fig. 1.**
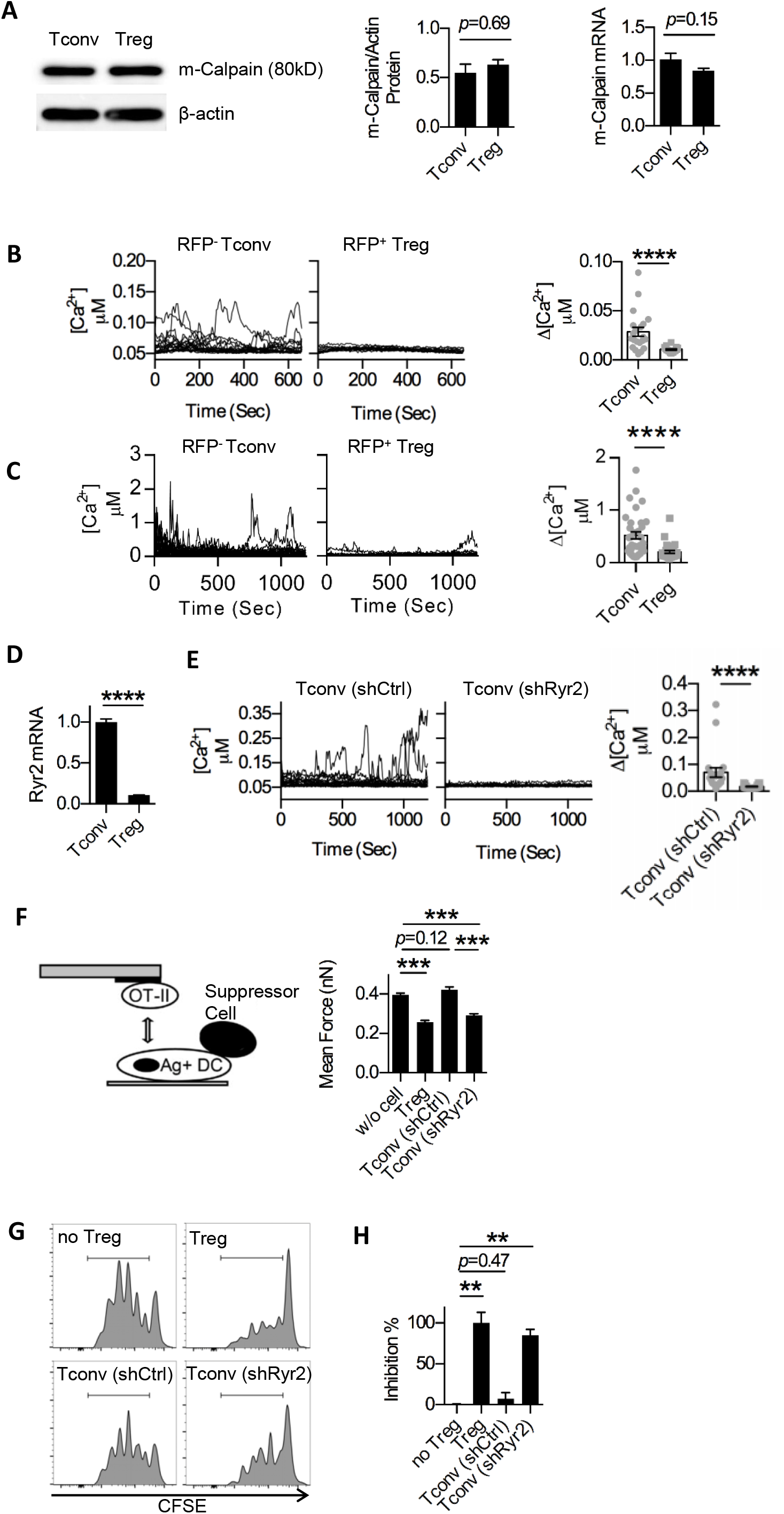
Reduced RyR2 activity is the basis of contact-dependent suppression. (**A**) Western blot protein (left) and qPCR RNA (right) analyses of m-calpain expression in Tconv and Treg cells isolated from FoxP3-GFP mice. Henceforth, all values are mean±SEM when applicable, N: number of independent experiments, N=4. (**B**) The change of intracellular free Ca^2+^ concentration [Ca^2+^] over time (left) and corresponding amplitude (right) were showed. Resting RFP^-^ Tconv and RFP^+^ Treg cell was sorted from CD4^+^ splenocytes of FoxP3-RFP mice by FACS following CD4 immunomagnetic negative selection (MACS) and loaded with Fluo-4 AM with 1.2 mM Ca^2+^ after overnight culture, respectively. One line represents one cell. Henceforth, n: number of biological repeats in one group of data points, n=20, N =5. **(C)** Same as B, with ratiometric Ca^2+^ imaging using Cal Red R525/650. n=20, N =3. **(D)** qPCR analysis of Ryr2 gene expression in FACS or MACS purified Tconvs and Tregs cultured overnight. N>5. (**E**) Same as B, shRyr2-treated Tconvs were used in place of Tregs. n=20, N=3. (**F**) Adhesion between OT-II T cells and OVA-pulsed DC2.4 cells that were free or engaged by Treg, RyR2 knockdown or control Tconv on the opposite side of the DC cell bodies was analyzed. Shown are the triple-cell AFM assay setup (left) and adhesion forces (right). N=3. (**G** and **H**) DC occupation by Treg, control or RyR2 knockdown Tconv cell-mediated suppression of OT-II T cell division. G: FACS plots, normalized to Mode; H: relative efficiency of inhibition using Treg and no Treg as 100% and 0%, respectively. N=3. Here, **, < 0.01; ***, < 0.001; ****, < 0.0001.

Upon T cell activation, Ca^2+^ signal comes from IP3R Ca^2+^ release channels that expands to secondary Ca^2+^ amplification events, such as the opening of store-operated Ca^2+^ channels (*17*). This activation-induced signal is absent in resting cells. To reveal why the spontaneous Ca^2+^ oscillations are lower in Tregs, we scanned typical calcium regulators by qPCR and found RyR2, an ER-membrane calcium channel protein, showing a markedly reduced expression in Tregs (fig. S1B and Fig. 1D). To see whether the reduction of RyR2 activities resulted in the increased binding to DCs, we performed SCFS along with several control Ca^2+^ channels. RyR2 reduction increased the T cell binding to dendritic cell (fig. S1B). To see if RyR2 function is indeed reduced in Tregs, we used its stimulator 4-CMC, and observed the rate of CMAC (calpain substrate) digestion. While 4-CMC induced calpain activities, Tregs were insensitive to this treatment, confirming the reduced expression of RyR2 (fig. S1C). JTV519, a RyR2 inhibitor, reduced Ca^2+^ levels in resting Tconvs to the level of Tregs, and elevated the binding force to DCs (fig. S1D). In knock-down assays, RyR2-specific shRNA reduced RyR2 mRNA level (fig. S1E). This was correlated by a reduction in the spontaneous Ca^2+^ oscillations in the treated Tconv cells (Fig. 1E), in line with reduced Calpain substrate digestion (fig. S1F).

To confirm that this increased binding excluded antigen-specific Tconvs to engage the same DCs, we performed the triple cell binding force analysis (*2*). Resembling Treg-mediated blockage as we reported previously, RyR2 KD Tconvs inhibited the binding strength between OTII and OVA-loaded DCs (Fig. 1F). Therefore, we confirmed at the single cell level that a Tconv with reduced RyR2 activity is functionally equivalent to a Treg in their ability to block Tconv-DC interaction. shRNA-treated Tconvs also suppressed OTII division in response to OVA pulsed DCs (Fig. 1, Gand H), confirming the unique involvement of RyR2.

### RyR2 is transcriptionally silenced by FoxP3

To confirm the exclusivity of RyR2 reduction in Tregs, it is essential to establish its link to FoxP3. We therefore overexpressed FoxP3 in T, A20, 3T3, and Renca cells. In all cases, the presence of FoxP3 reduced the level of RyR2 mRNA, suggesting that FoxP3 targeting of RyR2 is autonomous (Fig. 2A). Coincidentally, a previous report analyzing the binding of overexpressed FoxP3 in Tconvs produced a whole genome ChIP-Seq dataset for gene expression analysis (*18*). When we used IGV program to focus this dataset on a 40 kb region surrounding RyR2 gene, a signal in the 1.5 kb promoter region was detected over the untransfected control (Fig. 2B) (*19*). We cloned this region and constructed a luciferase reporter. To confirm the binding occurs in the native state, we performed ChIP QPCR analysis with WT Tregs (Fig. 2C). Although the signal ratio is not as strong as a FoxP3 overexpression system (fig. S2), FoxP3 showed a detectable binding to this region. Further truncation experiments narrowed the suppression activity to a defined region between ∼300-500 bp after the TSS site (∼200 bp before ATG). We then analyzed this refined segment for any potential FoxP3 binding sites. A GCAGGGG sequence, reported in a previous paper to be targeted by FoxP3, appears twice in the vicinity (*20*). When these two sites were depleted, the FoxP3 overexpression lost its ability to suppress luciferase, indicating site-specific FoxP3 binding (Fig. 2D). These data indicate that RyR2 is indeed under direct control of FoxP3.

**Fig. 2.**
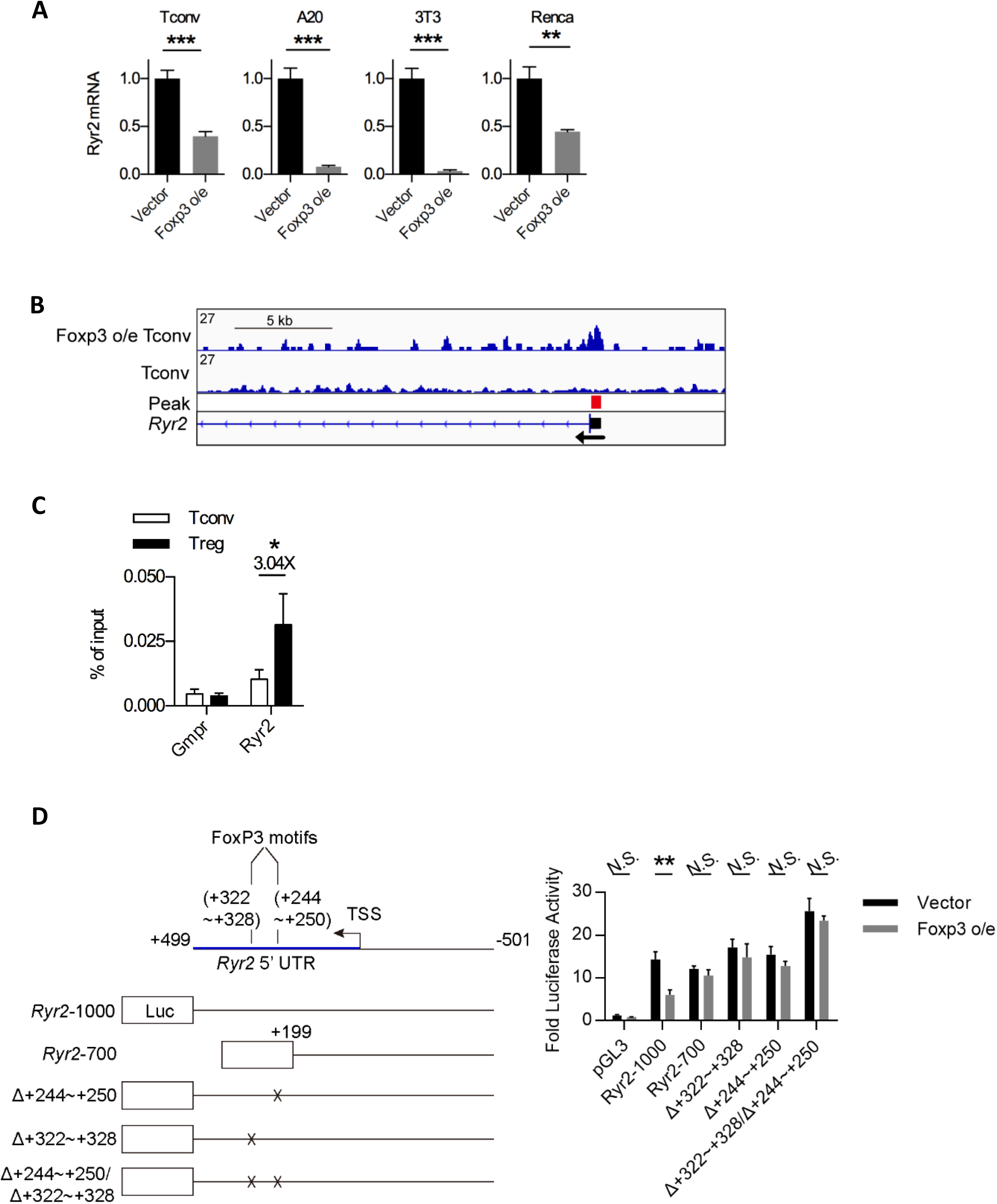
RyR2 is transcriptionally silenced by FoxP3. (**A**) RyR2 mRNA level in FoxP3-overexpressed Tconv, A20, 3T3 and Renca cells was detected by qPCR. N=3. (**B**) Anti-flag-FoxP3 ChIP-seq dataset re-analysis was performed in Tconvs with (upper track) or without (lower track) transduction of flag-FoxP3. Shown is the region around TSS (Transcription Start Site) of Ryr2 gene. Red box is the position of significant difference identified by the program, which mostly overlaps the promoter region (dark box) of this gene. (**C**) ChIP-qPCR analysis was performed in overnight cultured Tconvs and Tregs in presence of recombinant IL-2 to examine FoxP3 enriched binding in Ryr2 promotor region. Gmpr is negative control. N=3. (**D**) Left: Schematic diagram of the Ryr2 promoter-luciferase reporter constructs, with deleted positions indicted by ×. Right: Analysis of different truncated or FoxP3-binding sequence consensus-deleted Ryr2 promoter-driven transcription response in FoxP3-overexpressed 3T3 cells. TSS, Transcription Start Site. N=4. Here, *, < 0.05; **, < 0.01; ***, < 0.001; N.S., not significant.

### RyR2-deficiency minimally affects T cell development

Likely due to its exceptionally large size and complex regulation, thus far, overexpression of RyR2 has not been feasible in primary cells (*16*). We attempted to produce two mouse models with elevated RyR2 in Tregs: one with RyR2 under a strong viral promoter and the other with a FoxP3 conditional knock in where the two FoxP3 binding sites were mutated. Neither of them showed increased RyR2 in Tregs. Our attempt to drive the expression of RyR2 with VP64/dCas9 also failed (fig. S3A) (*21*). However, it was possible to delete RyR2 in CD4^+^ T cells with CD4-cre (fig. S3B). RyR2 deletion in Tconvs did not result in any overt change in mice, including the rate of development and body weight. Both thymic and peripheral CD4^+^ vs CD8^+^ marker distributions were nearly identical to WT mice (fig. S3C). The percentage of FoxP3^+^CD4^+^ T cells also remained undisturbed (fig. S3D). Other T cell activation indices were also similar including the frequency of CD44hi T cells, cytokine expression, and CD39/Helios/CD5 levels as measurement of self-reactivity (fig. S3E). The only difference was the percentage of Ki-67^hi^ CD4^+^ Tconvs, a phenomenon likely related to suppressive environment in the CKO mice (see later text). To further reveal any impact of RyR2 deletion in T cell biology, we used 50:50 WT/RyR2 CKO bone marrow to restore immune system in γ-irradiated recipients. Using CD45.1/CD45.2 markers, we repeated the thymic and peripheral comparisons (fig. S3F), and no difference was found. Therefore, the RyR2-specific deletion did not result in developmental defects in T cell subsets. This deletion was accompanied by reduced spontaneous Ca^2+^ oscillations in CD4^+^ T cells (Fig. 3A). In comparison to untreated Tconvs, CMAC digestion was reduced in RyR2^-/-^ Tconvs (fig. S3G). Similarly, force spectroscopy also indicated an increased binding to DCs upon RyR2 deletion (Fig. 3B). This was accompanied by their enhanced ability to interfere with the DC contact with T cells in the triple cell force analysis (Fig. 3C). More importantly, the RyR2^-/-^ Tconvs now behaved similar to Tregs in their ability to suppress OTII expansion stimulated by antigen-positive DCs (Fig. 3D).

**Fig. 3.**
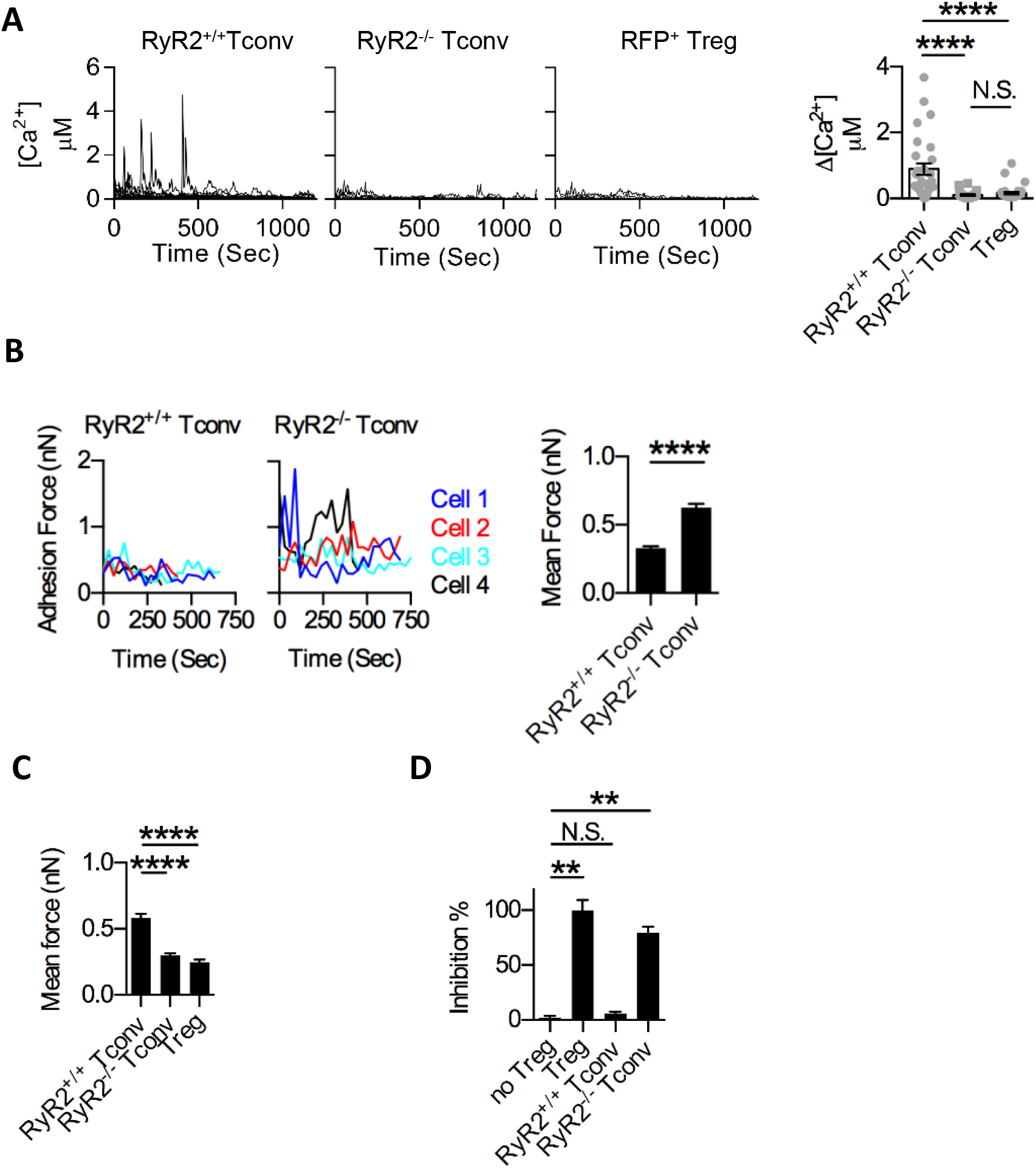
RyR2-deficiency minimally affects T cell development. (**A**) Resting RyR2^+/+^ Tconv, RyR2^-/-^ Tconv and Treg cells were loaded with Cal Red R525/650-AM and Ca^2+^ concentration fluctuations were analyzed. One line represents one cell (left). Corresponding amplitude was shown in the right. n=20, N =5. (**B**) SCFS force readings for RyR2^+/+^ Tconv and RyR2^-/-^ Tconv cell adhering to DC2.4 (left and middle) and their mean forces (right) were shown. N=3. (**C**) Mean adhesion forces between OT-II T cells and OVA-pulsed DC2.4 cells that were free or engaged by Treg, RyR2^+/+^ Tconv or RyR2^-/-^ Tconv cell on the opposite side of the DC cell bodies were shown. N=3. (**D**) RyR2^+/+^ or RyR2^-/-^ Tconv-mediated suppression of OT-II T cell division was analyzed. Shown was the relatively inhibition efficiencies of Tregs, RyR2^+/+^ and RyR2^-/-^ Tconvs. The efficiency of inhibition of Treg and no Treg were as 100% and 0%, respectively. N=3. Here, **, < 0.01; ****, < 0.0001; N.S., not significant.

### RyR2 deficiency per se mediates contact-dependent suppression

It remained possible that RyR2 deletion in Tconvs triggered another suppression mechanism other than our DC occupancy-based proposal. We found that RyR2^-/-^ cell surface markers potentially associated with Treg functions remained unchanged (fig. S4A). TGF-β and IL-10 production was undetectable (fig. S4B). To globally analyze the impact of RyR2 deletion, we performed RNAseq and ATACseq analyses for Tconvs, RyR2 CKO (RyR2^-/-^) Tconvs and Tregs, both under resting state and following activation. RNAseq revealed that with the expected reduction of RyR2 in both Tregs and CKO, no other suspected suppression function is significantly elevated in CKO (Fig. 4A). For the totality of data, we performed Spearman correlation and hierarchical clustering among RNAseq samples (fig. S4C). In summary, among the three cell types, the activation vs resting represented the first order (largest) of differences. The second order of differences was between Treg and Tconv/CKO. The last order was between Tconv and CKO. Therefore, CKO and Tconvs were similar, and both of them shared a farther distance to Tregs. ATACseq result showed a similar trend (Fig. 4B: FoxP3, helios and S1P receptor). Again, we failed to identify any major chromatin accessibility change in RyR2-deleted Tconv cells that resembled Tregs. Spearman correlation also showed the similarity between Tconvs and RyR2 CKO Tconvs, both as a group more distant from Tregs (fig. S4D). We also compared the expression level of Forkhead family genes and T cell associated calcium regulators in searching for any compensatory upregulation, no major change was found in RyR2 CKO cells (fig. S4, E and F). Those results seem to suggest that by simply limiting RyR2 expression, the Tconvs gained the ability to suppress DC-mediated T cell activation *in vitro*, without invoking another overt suppression program.

**Fig. 4.**
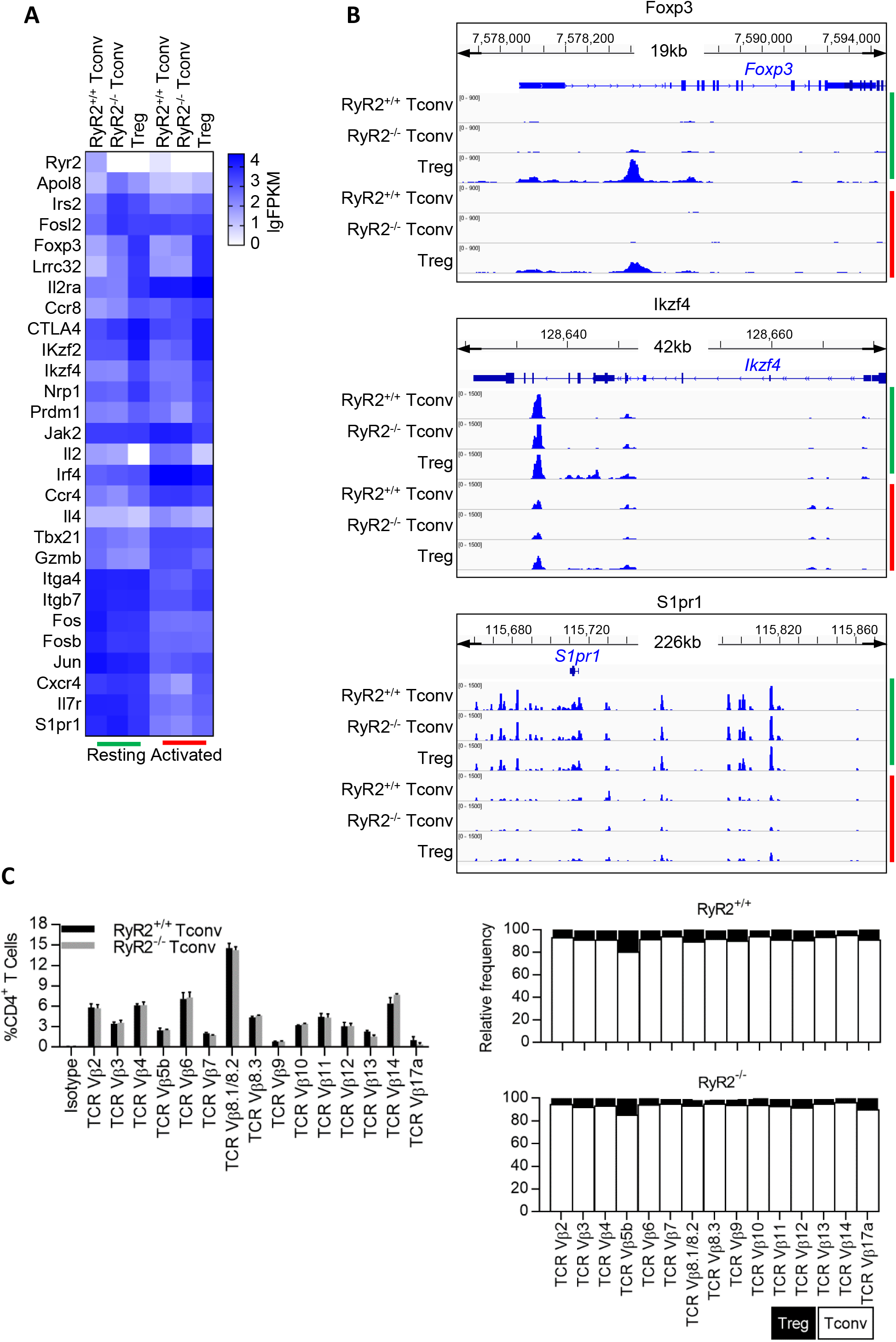
RyR2 deficiency per se mediates contact-dependent suppression. (**A**) Transcriptional difference among RyR2^+/+^, RyR2^-/-^ Tconv and Treg cells, before (green) and after anti-CD3/CD28 activation (red). Differential expressed genes were plotted by log10-FPKM. (**B**) Shown was chromatin opening status of representative genes in the three cells, before (green) and after activation (red). (**C**) T cell receptor Vβ chains of splenic CD4^+^ T cells from CKO and WT mice were assessed using a commercial mouse TCR Vβ screening panel by Flow cytometry. The expression of TCR vβ chains in splenic CD4^+^ T cells for peripheral TCR usage were analyzed. N=3.

### RyR2 deficiency-mediated suppression operates in the absence of specific antigen

A TCR repertoire analysis revealed that RyR2 CKO TCR frequencies were almost identical to WT Tconvs, and distant from Treg (Fig. 4C). Therefore, the suppression by RyR2 CKO Tconvs does not appear to involve significant change in T cell specificity. In our previous reports, we found that Tregs showed prolonged contact with DCs *in vivo* (*7*). Another report, mostly using induced, antigen-specific iTregs, suggested that iTreg suppression intensity was related to antigen specificity (*9*). We performed the intra-vital imaging assay with RyR2 CKO Tconvs. Fig. S5A shows that CKO Tconvs exhibited an extended contact with DCs *in vivo*, similar to Tregs. To further reveal if TCR specificity contributed to the extended binding, we also produced RyR2 CKO OT II T cells and repeated the experiment. Infusion of OVA peptide increased the binding duration between WT OTII T cells and DCs, yet it had no effect on contact duration between RyR2 CKO OTII T cells and DCs, suggesting the adhesion to DCs can be inversely regulated by RyR2, and the antigen specificity may not be essential in our settings (fig. S5B). To calculate the impact of CKO/Treg binding on DC’s ability to interact with antigen-specific responder T cells, we infused OT II T cells with or without antigen, in the presence of either control Tconvs, Treg or CKO Tconvs. The antigen increased the contact time between DCs and responder OT II cells; whether DC was bound by a control Tconv or not did not change any contact duration for the former pair. However, if the DC was bound by Treg or CKO, the DC/responder OT II contact time was significantly reduced (Fig. 5A). Ca^2+^ reporter activities in the responder OT II cells translated the contact duration differences into T cell activation intensity (Fig. 5B). To address the issue of Treg/CKO antigen specificity, we used this 3-cell system again in the constant presence of OVA. In this system, OT II Treg and WT Treg showed identical ability to block DC/responder OT II contact, and the same results were obtained from CKO and OT II CKO, suggesting that antigen specificity of tTreg or CKO did not affect their suppression efficacy (Fig. 5, C and D). Ca^2+^ reporter activities in responder OT II cells were consistent with those data (Fig. 5, C and D). Therefore in our system, antigen specificity of Treg and CKO Tconv itself does not appear to alter their ability to suppress DC’s engagement to other T cells.

**Fig. 5.**
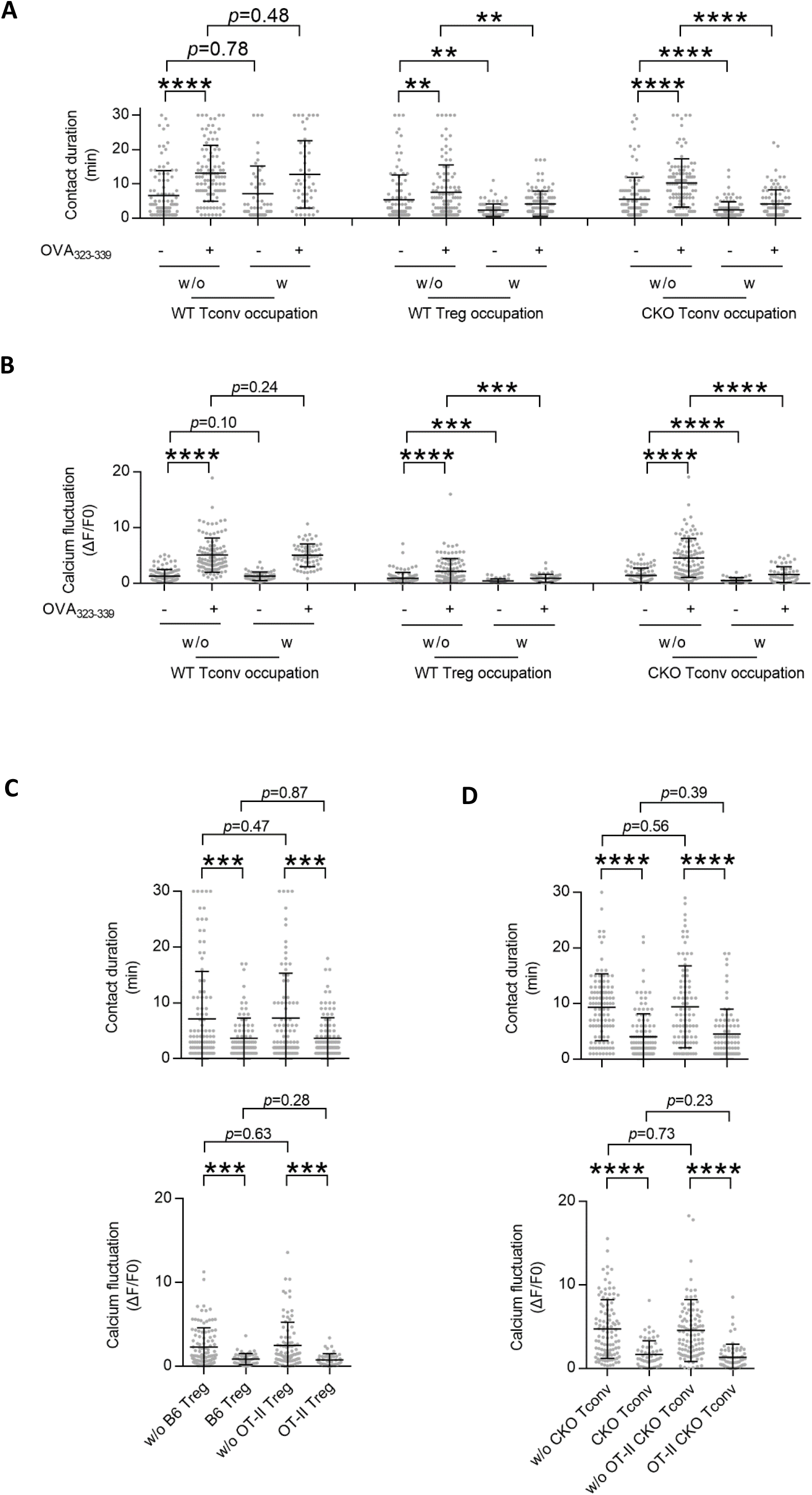
RyR2 deficiency-mediated suppression operates in the absence of specific antigen. (**A**) Contact disruption of responder OT-II and DCs by suppressor cells in intra-vital imaging. Wildtype Treg and RyR2^-/-^ Tconv cells (CKO) were analyzed as suppressor cells, whereas wildtype Tconv cells were added as negative control. Both antigen non-specific and OVA_323-339_ specific contacts were analyzed. 100 contacts were analyzed for each group. w/o, OT-II-DC contacts without suppressor cell occupancy. w, OT-II-DC contacts with suppressor cells on the specific DC. n=4, N=2. (**B**) Calcium signal of OT-II T cells during contacts. OT-II cells preloaded with calcium indicator FuraRed were imaged and analyzed for each condition. 100 cells were analyzed for each group. n=4, N=2. (**C** and **D**) Contact disruption of OT-II cells and antigen-loaded DCs by suppressor cells. Wildtype Treg (non-specific) and OT-II Treg (specific) were analyzed in C. RyR2^-/-^ Tconv (non-specific) and OT-II-RyR2^-/-^ Tconv (specific) were analyzed in D. 100 contacts were analyzed for each group. w/o, OT-II-DC contacts without suppressor cell occupancy. w, OT-II-DC contacts with suppressor cell on the specific DC. 100 cells were analyzed for each group. n=4, N=2. Here, **, < 0.01; ***, < 0.001; ****, < 0.0001.

### RyR2-deficient Tconvs are indistinguishable from Tregs in disease models and scurfy rescue

To establish functional equivalency of RyR2^-/-^ Tconvs and Tregs, we tested their behaviors in viral infection, allergic response, autoimmune colitis and tumor development, with comparison to similarly infused purified Tregs. In a footpad inoculation model, HSV-1 pfu counts were increased by about roughly one log with either Tregs or RyR2^-/-^ Tconvs. Control infusion of Tconvs showed no increase over the viral inoculation alone (Fig. 6A) (*22, 23*). With a secondary challenge, this same model has been used to demonstrate DTH response (*24*). Sensitized footpads were swollen upon 2^nd^ HSV-1 inoculation, yet the increase in the thickness was reduced by both Tregs and RyR2^-/-^ Tconvs, but not by control Tconvs (Fig. 6B, image: fig. S6A). In an OVA sensitization-induced asthma model, infusion of Tregs and RyR2^-/-^ Tconvs were equally effective in limiting BALF cell number counts, with reductions to a similar degree in both lymphocytes and eosinophils (Fig. 6C, sensitization schedule and histology: fig. S6B). In DSS-induced colitis model, the colon length was reduced. Both Tregs and RyR2^-/-^ Tconvs, but not control Tconvs, were able to reverse the reduction and limit the colon damage (Fig. 6D, induction schedule, colon images and histology: fig. S6C).

**Fig. 6.**
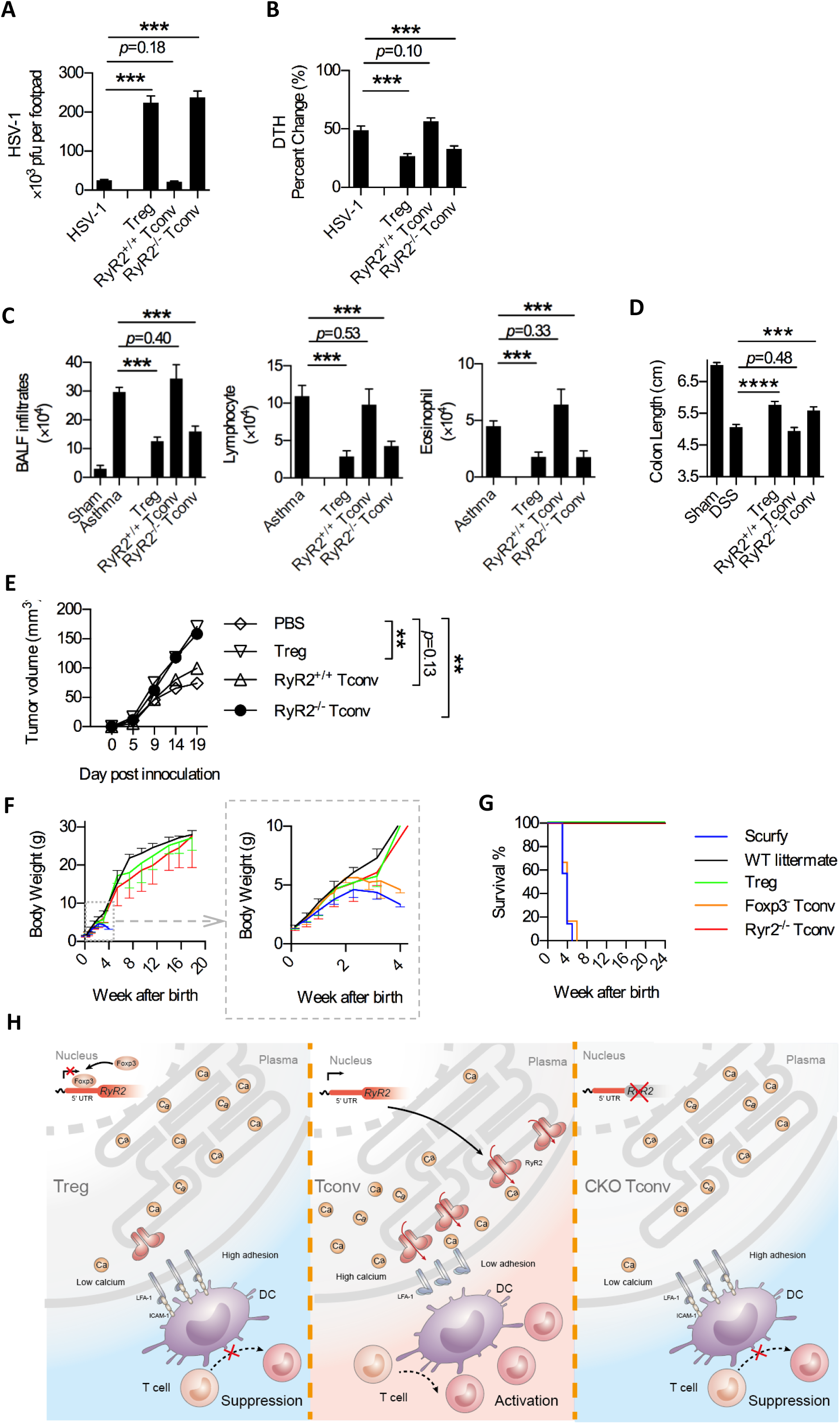
RyR2-deficient Tconvs are indistinguishable from Tregs in disease models and scurfy rescue. (**A**) HSV-1 footpad infection model. As described in the methods, virus was infected at day 0, 2×10^5^ cells were transferred into footpad at day 3. At day 7, analysis of HSV-1 titer in footpad tissues was done. n=4-6. N=3. (**B**) After UV-inactivated HSV-1 antigen re-challenge in footpad at day 6, analysis of delayed-type hypersensitivity (DTH). Footpad thickness without re-challenge was set as 0%. Data are pooled with a total n=16-21 mice per group. (**C**) OVA-induced asthma model. As described in methods, mice were sensitized at day 0 and day 14, intra-tracheal re-challenged at day 21, 23 and 25. 10^6^ cells were transferred i.v. at day 23. At day 32, analysis of total infiltrates, infiltrated lymphocytes and eosinophils in BALF. N=3. (**D**) DSS-induced colitis model. After adoptive transfer of 3×10^6^ Treg, RyR2^+/+^ or RyR2^-/-^ Tconv cells on day -1, DSS water was provided from day 0 to day 7. Colon length was measured on day 9 in sham or DSS-treated mice. Pooled data, n = 11-18 mice/group. (**E**) MC38 tumor model. Tumor growth of C57BL/6 mice inoculated with MC38, MC38 plus 10^6^ Treg cells, RyR2^+/+^ or RyR2^-/-^ Tconv cells. Shown were pooled caliper measurements. Pooled data, n = 10 mice per group. (**F** and **G**) Rescue of Scurfy mice. (F) Analysis of weight change over time following i.p. transfer of Treg, FoxP3^-^ Tconv or CKO Tconv cells into newborn Scurfy mice. n=10 for WT littermate, n=7 for Scurfy, n=6 for Treg, n=6 for FoxP3^-^ RyR2^+/+^ Tconv, n=6 for RyR2^-/-^ Tconvs. Full 20-week rescue was shown in the left, and 4-week weight change of Scurfy and FoxP3^-^ RyR2^+/+^ Tconv was shown in the right. (G) Kaplan-Meier survival curves for the mice described in F. (**H**). Proposed working mechanism: In Treg cells, FoxP3 expression autonomously suppresses the expression of RyR2, targeting a stretch of sequence roughly 200 bp before RyR2 start codon, which results in severely depressed basal calcium oscillation in Tregs. The depressed basal Ca^2+^ level is insufficient to activate m-Calpain to cleave LFA-1-anchoring proteins, such as talin, when Treg cells contact with DCs. This causes Tregs to adhere to DCs with exceedingly strong force, rendering the latter incapable of engaging other Tconvs. This mechanism is independent of other immune regulatory functions, under direct control of FoxP3 and sufficient to explain Treg effector functions in most disease models and in Scurfy mice. Here, **, < 0.01; ***, < 0.001; ****, < 0.0001.

Finally, in MC38 tumor model (*25*), the infusion of both Tregs and RyR2^-/-^ Tconvs facilitated the tumor growth, with control Tconvs showing no overt effects (Fig. 6E). Therefore, in several disease models whereby Tregs are known to show immune regulatory roles, those effects are phenotypically copied by RyR2^-/-^ Tconvs in the absence of FoxP3. The lack of functional Tregs is most evident in FoxP3-deficient mice (or Scurfy) with pervasive inflammation as a result of multi-organ autoimmunity. This pathology leads to death in a defined window of 2-4 weeks of age. In addition to the immune regulatory effects of RyR2^-/-^ Tconvs, if this regulation is also the effector function missing in Scurfy mice, RyR2^-/-^ Tconvs should correct those autoimmunities in the systemic absence of FoxP3. Scurfy mice (FoxP3^-/-^, C57BL/6) were injected with PBS, Tregs, FoxP3^-^ Tconvs or RyR2^-/-^ Tconvs 2-3 days after birth. As expected, mice infused with PBS or FoxP3^-^ Tconvs all died within a window of 2-4 weeks after birth (Fig. 6F, G). In contrast, all those infused with Tregs or RyR2^-/-^ Tconvs had survived more than one year without any sign of shortened life expectancy. For the histology, control-infused mice were analyzed on week 3, and those from Treg and RyR2^-/-^ Tconv-infused were on week 8-12. As expected, PBS or FoxP3^-^ Tconv-infused mice showed severe thyroiditis, splenitis, pneumonitis, dermatitis, hepatitis, pancreatitis, gastritis, and colitis (fig. S6D). The infusion of either Tregs or RyR2^-/-^ Tconvs prevented those pathologies, providing evidence that the regulation of RyR2 is a potentially generalizable effector mechanism of Tregs (Fig. 6H).

## DISCUSSION

FoxP3 being the master regulator of Tregs is built on the observation that its expression in T cells endows the suppressive phenotype (*26, 27*). The finesse of this complex regulation on this one transcriptional factor is being appreciated at ever-deepening levels. Recent advances on additional regulations, such as via Bcl11b, CDK8/19, and FoxP1 introduce additional complexity (*28-30*). Even its downstream factors, such as BLIMP1, has been a new focal point as it fine-tunes Tregs in adipose tissues by sex-specific factors (*31*).

In comparison to its gene regulation, the study on Treg effector mechanisms has been more diverse. Trogocytosis of costimulatory molecules on DCs via CTLA4 expressed on Tregs, MHC class II molecule depletion on DCs are two leading hypotheses. Those topological extractions, along with previously reported suppressive cytokines, metabolic blockage, direct T cell cytolysis, IL-2 deprivation etc, were all effective in respective settings (*1*). While those important advancements build our understanding of Tregs, we still cannot come to an unequivocal conclusion whether there is a specific suppression mechanism. Conceptually, the suppressive capacity is bestowed by FoxP3 expression, the question becomes whether such a switch turns on something fundamentally unique to Tregs. As existing proposed mechanisms are not exclusively controlled by FoxP3, a “truce” can be reached that all these mechanisms are working together to create a network of suppression. However, this synthesis is not ideal in that the complexity to resolve the details of such a FoxP3-originated cooperation is almost experimentally unattainable. Empirically, evidence of those factors working together has been lacking in disease settings or animal models.

We believe that other suppression mechanisms, i.e. biophysical and spatial-temporal, may be at work as Tregs’ effector function, particularly while the strong binding of Tregs with DCs is an observation that has been repeatedly made by many labs in the field. Our labs proposed in 2017 that spatial occupation of DCs by Tregs limits the former’s ability to engage other T cells (*2, 7*). Evidence was provided in the form of *in vivo* contact inhibition and force spectroscopy analysis that the binding essentially blocked the DCs from forming steady contact with Tconvs. We also provided evidence that binding was due to the limited m-calpain activities in Tregs (*2*). However, why Tregs have limited m-Calpain activity was unknown.

In immune cells, the most studied Ca^2+^ activation is initiated by IP3, being one of the key downstream events of ITAM phosphorylation-mediated enzymatic functions common to FCR, TCR and BCR. The release of soluble inositol triphosphate by PLC cleavage triggers the opening of IP3R on the ER membrane. This strong Ca^2+^ flux is receptor ligation-dependent and unique in immune cells. RyR family members, best studied for the control excitation–contraction coupling in muscle cells and synaptic transmission, are ubiquitously expressed and involved in membrane-localized cytoplasmic Ca^2+^ homeostasis (*13*). Similar to IP3 signals, those events can be trigger-dependent (i.e. membrane depolarization). However, RyRs also control the basal Ca^2+^ oscillation (*32*) and those Ca^2+^ activities are gradually coming into our focus as being essential to basic cell biology (*33*). Importantly, Ca^2+^ signaling is likely the most complex regulation in biological systems, with variations in pattern, duration, and subcellular location, and is not mere a reflection of intensity. A distinct Ca^2+^ signal often reaches its intended target while multiple other Ca^2+^ events are concurrent (*34*). In T cells, IP3R and SOCE (store-operated calcium entry) channels are the dominant signaling during activation, with the latter providing the “global” Ca^2+^ waves that can reverberate for minutes to hours. This is in sharp contrast with the “puff”-like, PM inner leaflet-targeting RyR2 signal that precisely localized at PM (plasma membrane)-ER junction (*33*). As m-Calpain is active only in its membrane associated state, this association may isolate the RyR2-m-Calpain regulation from the rest of T cell calcium events.

In this report, we found that Tregs, in comparison to other T cells, lack the basal Ca^2+^ oscillation, a phenotype as a result of severely suppressed RyR2 expression. This baseline Ca^2+^ flux is required for m-Calpain activities. Suppression of RyR2 channel activities or genetic blockage of its expression renders Tconvs to behave as Tregs in their suppression. The most critical finding is that RyR2 expression is directly blocked by FoxP3 expression irrespective of cell types, demonstrating a true causative autonomy. RyR2 conditional deletion results in the entire pool of Tconvs to become suppressive with immune inhibitory capacities indistinguishable from those of Tregs (Fig. 6H). Perhaps more tellingly, those Tconvs correct FoxP3 deletion-associated Scurfy phenotype and restore immune homeostasis in the absence of FoxP3. In our survey of RyR2^-/-^ Tconvs, factors that may be associated with immune suppression do not appear to be altered, suggesting that this regulation may be independent of previously proposed models. This certainly does not exclude those factors from participating in Treg suppression in other disease models. Yet, the RyR2 deletion resulting in an immune suppression similar to Tregs suggest that this chain of regulation may be one of the central effector mechanisms of Tregs, fulfilling a missing link in this research field.

Another point of potential interest is the implication of our work on the debate of whether Tregs must be antigenic-specific to exert their immune suppression. For the Treg pool, at least tTregs are believed to originate from the thymus as a group of CD4^+^ T cells carrying highly self-reactive TCRs. Tconv TCRs, on the other hand, are shaped more stringently by the thymic deletion and are therefore unlikely to be the same as those on tTregs (*35*). As RyR2^-/-^ Tconvs are also potently immune suppressive, similar to Tregs, our results seem to suggest at the moment of suppression, a strong signal via Treg TCR may not be essential, which explains the equal inhibitory potency between antigen specific and non-specific Tregs/RyR2^-/-^ Tconvs in our *in vivo* imaging results. This notion certainly does not rule out that in certain settings, particularly those induced Tregs rising in inflammatory milieu, antigen specificity may be quantitatively advantageous, thus providing two layers of suppression to better suit the varying degrees of inflammatory attacks on the host.

## MATERIALS AND METHODS

### Mice

All mice were in CD45.2 C57BL/6 background unless noted otherwise. RyR2^fl/fl^ mice were made in GemPharmatech Co.Ltd and verified by genotyping. OT-II-transgenic mice and CD45.1 mice were purchased from Jackson Laboratories. CD11c-DTR-eGFP transgenic mice were a gift from Dr. Yonghui Zhang, School of Pharmaceutical Sciences, Tsinghua University. Female FoxP3^+/-^ mice and FoxP3-RFP mice were a gift from Dr. Xuyu Zhou, Institute of Microbiology, Chinese Academy of Sciences (CAS). FoxP3-IRES-GFP and Foxp3-Cre-YFP transgenic mice were a gift from Dr. Hai Qi, School of Medicine, Tsinghua University. CD4-Cre mice were a gift from Dr. Chen Dong, School of Medicine, Tsinghua University. Wild Type mice were purchased from Beijing Vital River Laboratory Animal Technology Co., Ltd. All mice were bred and housed at Tsinghua University Animal Facilities and maintained under specific pathogen-free conditions. All animal experiments were conducted in accordance of governmental and institutional guidelines for animal welfare and approved by the IACUC at the Tsinghua University.

### Cell Lines and Primary Cell Culture

DC2.4 Cells were a gift from Dr. Ken Rock of UMass Medical School. Vero cells were from Dr. Xu Tan of School of Pharmaceutical Sciences, Tsinghua University. Renca cells were from Dr. Guangyu An, Chao-Yang Hospital, Beijing. HEK293FT cells were a gift from Dr. Wei Guo of School of Medicine, Tsinghua University. MC38 and NIH-3T3 cells were purchased from the American Type Culture Collection (ATCC). MC38, Renca, NIH-3T3 cells and Vero cells were cultured in Dulbecco’s modified Eagle’s medium (DMEM) containing 10% FBS, 100 U/ml penicillin, 100 ug/ml streptomycin. All other cells were grown in RPMI-1640 with the same supplements plus 10 mM Hepes (pH 7.0) and 50 μM β-mercaptoethanol. All cell lines were tested for mycoplasma contamination by PCR analysis. Murine CD4^+^CD25^+^ Tregs and CD4^+^CD25^−^ Tconvs were isolated from spleens using mouse CD4^+^ T Cell Isolation Kit (StemCell, 19852) and mouse CD25 Regulatory T Cell positive selection Kit (StemCell, 18782). Treg and Tconv cells sometimes were sorted by FACS from CD4^+^ splenocytes of Foxp3-IERS-GFP or Foxp3-IERS-RFP transgenic mice. Murine DCs were isolated with mouse CD11c selection Kit II (StemCell, 28007) from spleens. OT-II T cells were isolated from OT-II splenocytes by mouse CD4^+^ T Cell Isolation Kit (StemCell, 19852) and sometimes sorted by FACS with an anti-TCR Vα2 antibody (eBioscience, B20.1).

### Antibody and Reagents

Recombinant Human IL-2 was from R&D systems. Dual-Luciferase Report Assay System was purchased from Promega. Anti-mouse CD3ε monoclonal (145-2C11), anti-mouse CD28 monoclonal (37.51) and anti-mouse FoxP3 monoclonal antibodies were from eBioscience. For flow cytometric analysis, in addition to CD45.1 (A20) monoclonal antibody from Invitrogen, others were purchased from eBioscience or BD Pharmingen. The following clones were used: CD45.2 (104), CD3 (17A2), CD4 (GK1.5), CD8a (53-6.7), FoxP3 (3G3), GITR (DTA-1), CD25 (PC61.5), PD-1 (J43), CTLA-4 (UC10-4B9), CD39 (24DMS1), Tim3 (RMT3-23), LAG-3 (C9B7W), TCR Vα2 (B20.1), CD44 (IM7), CD62L (MEL-14), CD5 (53-7.3), Nrp1(3DS304M), Helios (22F6), IFN-γ(XMG1.2), IL-4 (11B11), IL-17A (eBio17B7), Ki-67 (SolA15) and Rat IgG2a kappa Isotype Control (eBR2a).

### Western Blot

The cells were collected and lysed with RIPA buffer (Beyotime, P0013B). The cell lysate was centrifuged and the supernatant was collected. Total proteins were quantified with BCA Protein Assay Kit (Beyotime, P0012). After being mixed with 3×SDS loading buffer and boiled for 5 min, the proteins were loaded onto 7.5% or 5% PAGE Gels (EpiZyme, PG111). Then the proteins were transferred onto a Nitrocellulose membrane and immunoblotted with indicated primary (Anti-Ryanodine Receptor Antibody (C3-33), Thermo Scientific, 1:1000 dilution; anti-β-actin antibody (2D4H5), Protientech, 1:20000 dilution; m-Calpain large subunit (M-type) antibody, CST, 1:1000 dilution) and secondary antibodies (Anti-mouse IgG, HRP-linked Antibody, CST, 1:5000 dilution). Finally, the immunostained bands were detected by the Super ECL Detection Reagent (Yeasen, 36208ES76).

### Real-Time qPCR

Tconvs and Tregs were cultured overnight in presence of recombinant human IL-2 for subsequent experiments unless noted otherwise. Total RNA was extracted from indicated cells using TRIzol reagent (Invitrogen) and first strand cDNA was synthesized with Reverse Transcriptase M-MLV (TaKaRa). Real-time PCR was performed using Hieff® qPCR SYBR Green Master Mix (No Rox) (Yeasen). *gapdh* or 18S RNA was used as reference gene for normalization. The primer sequences were as follows: *gapdh*, 5’-CATCACTGCCACCCAGAAGACTG-3’ and 5’-ATGCCAGTGAGCTTCCCGTTCAG-3’ *18S RNA*, 5’-CGGACAGGATTGACAGATTG-3’ and 5’-CAAATCGCTCCACCAACTAA-3’ *Ryr2*, 5’-ATGGCTTTAAGGCACAGCG-3’ and 5’-CAGAGCCCGAATCATCCAGC-3’; *Ryr1*, 5’-GCACACTGGTCAGGAGTCGTATG-3’ and 5’-GGGTGTAGCACAGGATTTAT-3’; *Ryr3*, 5’-ATCGCTGAACTCCTGGGTTTG-3’ and 5’-TTCATGTCGATGGAACTTAGCC-3’; *FoxP3*, 5’-CCCATCCCCAGGAGTCTTG-3’ and 5’-ACCATGACTAGGGGCACTGTA-3’; *Capn2*, 5’-GGTCGCATGAGAGAGCCATC-3’ and 5’-CCCCGAGTTTTGCTGGAGTA-3’; *Itpr1*, 5’-CGTTTTGAGTTTGAAGGCGTTT-3’ and 5’-CATCTTGCGCCAATTCCCG-3’; *Trpm1*, 5’-ATCCGAGTCTCCTACGACACC-3’ and 5’-CAGTTTGGACTGCATCTCGAA-3’; *Trpm4*, 5’-GGACTGCACACAGGCATTG-3’ and 5’-GTACCTTGCGGGGAATGAGC-3’; *Trpv2*, 5’-TGCTGAGGTGAACAAAGGAAAG-3’and 5’-TCAAACCGATTTGGGTCCTGT-3’; *Cacna2d4*, 5’-GGCAGCAAGTTATCTCCCAG-3’ and 5’-CCACAGGATGATTGGCGTCTT-3’; *Ahnak*, 5’-CAGCGCATCTACACCACGAA-3’ and 5’-CACTTCATGCCTTGGTATCTTGA-3’; *Stim1*, 5’-TGACAGGGACTGTACTGAAGATG-3’ and 5’-TATGCCGAGTCAAGAGAGGAG-3’ *Pkd1*, 5’-CTAGACCTGTCCCACAACCTA -3’ and 5’-GCAAACACGCCTTCTTCTAATGT -3’; *Ncs1*, 5’-AGCAAGTTGAAGCCTGAAGTT-3’ and 5’-GCTGGGGCAGTCCTTAATGAA-3’; *Slc24a3*, 5’-AGCAAGTTGAAGCCTGAAGTT-3’ and 5’-GCTGGGGCAGTCCTTAATGAA-3’; *Hspa2*, 5’-GCGTGGGGGTATTCCAACAT-3’ and 5’-TGAGACGCTCGGTGTCAGT-3’; *P2rx7*, 5’-GACAAACAAAGTCACCCGGAT-3’ and 5’-CGCTCACCAAAGCAAAGCTAAT-3’; *Itgav*, 5’-CCGTGGACTTCTTCGAGCC-3’ and 5’-CTGTTGAATCAAACTCAATGGGC-3’; *Ccdc109b*, 5’-CCACACCCCAGGTTTTATGTATG-3’ and 5’-ATGGCAGAGTGAGGGTTACCA-3’; *Pln*, 5’-AAAGTGCAATACCTCACTCGC-3’ and 5’-GGCATTTCAATAGTGGAGGCTC-3’; *Gsto1*, 5’-ATCCGGCACGAAGTCATCAAC-3’ and 5’-TGACAGATTCGGTGACCAAGT-3’; *Itpr2*, 5’-CTGTTCTTCTTCATCGTCATCATCATCG-3’ and 5’-GAAACCAGTCCAAATTCTTC TCCGTGA-3’; *Itpr3*, 5’-CTTTATCGTCATCATCATCGTGTTG-3’ and 5’-AGGTTCTTGTTCTTGATCATCTGAG CCA-3’.

### FoxP3 ChIP-qPCR

ChIP was conducted following the manufacturer’s instructions of SimpleChIP® Plus Sonication Chromatin IP Kit (CST. 56383). Overnight cultured-Treg and Tconv cells (approximately 2×10^7^ cells per assay) were fixed for 15 min at room temperature with 1% formaldehyde, then was digested with 45 cycles of sonication (Bioruptor). FoxP3 mAb (eBioscience, clone 150D/E4) or control rabbit IgG antibody (CST) were added to bind. Binding of Ryr2 promoter was determined by real-time quantitative PCR. The following primer pairs were used: *Gmpr* promoter,5’-CAGCTGGAACAGCCTTGGAA-3’, and 5’-AAATGTCAAGGCCCCTGTGA-3’; Il2ra promoter,5’-GGGTCAGGCCAACTTAGATGAG-3’, and 5’-CTCAACAAAGACTGAGAAGCAAGGT-3’; *Ryr2* promoter, 5’-TGCAGGGGGACCGACC-3’, and 5’-GTCACTGCTAACCAGGATGTTCTA-3’.

### Calcium Imaging

Overnight cultured-T cells were stained with 2 μM fluo-4 AM (Thermo Fisher) in Hank’s solution (SL6080, Coolaber) at 37 °C for 30 min. After washing and incubation, the cells were allowed to adhere to a poly-L-lysine (0.1 mg/mL; Sigma-Aldrich)-coated round glass slip mounted in a sandwiched, self-made chamber at room temperature. Excess non-adherent cells were removed by flushing with Hank’s solution after 15 min. Then the measurement chamber was placed on an Olympus IX-73 microscope equipped with a 20× (numerical aperture: 0.8) or 40× (numerical aperture: 1.2) Olympus objective. Fluorescent Ca^2+^ signals were recorded as a time lapse for 20 min with an interval of 6 seconds. The emission signals at 468–550 nm excited by 488 nm laser were recorded with a charge-coupled device camera (ORCA-AG, Hamamatsu). Data collection was controlled by NIS-Elements 3.0 software (Nikon). The mean fluorescence intensity changes over time for individual cells were analyzed by ImageJ and normalized to resting fluorescence F_0_ (Fluo-4 F/F_0_) after subtracting backgroud. Calcium concentration was analyzed based on following equation for fluo-4(*36*),

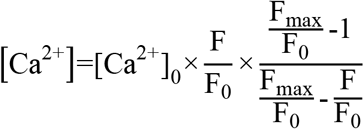

where Fmax is the fluorescence intensity at 10 mM saturating [Ca^2+^], the cytosolic [Ca^2+^]_0_ is 50 nM for resting T cells(*37*), respectively.

Ratiometric Ca^2+^ imaging was performed as described previously(*16*). CD4^+^ T cells were loaded with 5 μM Cal Red™ R525/650 AM (AAT Bioquest) at room temperature for 30 min. The dye-loaded cells were then subjected to wide-field imaging. The Cal Red™ R525/650 fluorescence dye was excited at 488 nm, the emitted fluorescence signal was captured at 525 nm (F525) and 650 nm (F650). For Cal Red, based on the following equation and known dissociation constants (Kd), the intensity ratios can be converted to intracellular calcium concentration(*38*).

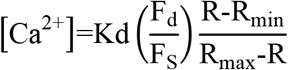

Where, R represents the fluorescence intensity ratio F525/F650 during the experiment, in which F525 and F650 are the fluorescence detection wavelengths for the ion-bound and ion-free indicator, respectively. R_max_ represents the ion-saturated fluorescence intensity ratio and R_min_ represents the completely ion-free fluorescence intensity ratio. F_d_/F_s_ is the ratio F_max_/F_min_ at the wavelength of the Ca^2+^-free form of the indicator (650 nm).

### Calpain Activity Measurement

For measuring calpain activities, 10^5^ overnight cultured-T cells were incubated at 200 μL PBS solution containing 20 uM calpain substrate CMAC (t-BOC-Leu-Met, Thermo Fisher) at room temperature in dark. Reaction was terminated by 4% PFA (Biosharp) up to 2 minutes after a 5-min or 1-min (for 4-CmC addition) incubation, then the cells were immediately placed on the ice at least 5 minutes. Calpain activities as fluorescence signals from the digested substrate were determined by Fortessa cytometers (BD Biosciences) via the Hoechst Blue channel. JTV519 was added 30 min before the experiment and 4-CMC was added at the start of assay.

### Flow Cytometry

For surface marker detection, cells were incubated with Fc blocker (CD16/32 antibody; 2.4G2) for 5 min, and then incubated with surface antibody for 15 min at room temperature avoiding light. Stained cells were analyzed directly or, occasionally, after being fixed with 1% paraformaldehyde (PFA) using Fortessa cytometers (BD Biosciences) with FACS Diva software. The FoxP3/Transcription Factor Staining Buffer Set (Invitrogen, 00-5523-00) was used for intracellular staining of mouse FoxP3, Ki67 and helios. The Intracellular Fixation/ Permeabilization Buffer Set (Invitrogen, 88-8824-00) was utilized for mouse cytokine detection. Data were analyzed with Flowjo V10.

### ELISA

For IL-10 and TGF-β detection, 10^6^ purified Tregs, RyR2^+/+^ and RyR2^-/-^ Tconvs were stimulated by anti-mouse CD3ε plus anti-mouse CD28 antibodies. After 72 hrs, the supernatants were collected. High-binding 96-well ELISA plates (Nunc) were coated with anti-mouse IL-10 (eBioscience) and anti-human/mouse TGF-β capture antibody (eBioscience) at 4°C overnight, respectively. After drying, plates were blocked with 2% BSA in PBS for 1 h at room temperature. After washing, 100 uL of diluted cell supernatant was added in triplicate wells, followed by incubation for 1 h at room temperature. Plates were then washed with PBST (0.05% Tween20, Sigma-Aldrich, in PBS), and were incubated with anti-mouse IL-10 and anti-human/mouse TGF-β detection antibody for 0.5 h at room temperature, respectively. TMB (eBioscience) was added (100 mL/well) and the plates were incubated for 10 min at room temperature in dark, followed by the addition of H_2_SO4 (50 uL, 1 M) per well to terminate the reaction. Optical density (OD) was immediately read at 450 nm using an ELISA plate reader (Bio-Rad).

### Gene Knockdown and Overexpression

Lentivirus-based shRNA was used to knock down specific genes. pLKO.1 vector purchased from shRNA library platform of Center of Biomedical Analysis of Tsinghua University were used for our all gene knockdown experiments. All pre-designed shRNA sequences were synthesized by Ruibiotech. Then they were inserted into pLKO.1 vector. Plasmids including shRNA and the package constructs (pMD2.G and psPAX2) were purified from transformed E. coli with EndoFree Plasmid Midi Kit (CWBIO, CW2105S). Lentivirus production was performed according to the manufacturer’s instruction. In brief, 293FT cells were cultured in a 10 cm dish with 60-80% confluency. Culture media were replaced 2 hrs before DNA transfer. 5 μg pLKO.1 vector (with inserted shRNA) was transfected into 293FT cells with 2.5 μg package vectors pMD2.G and 2.5 μg psPAX2 using Neofect™ DNA transfection reagent (Neofect, TF201201). 72 hrs later, lentiviruses were harvested and were added to infect T cells with Polybrene (final concentration 4 μg/ml). 48 hrs after the virus infection, cells were sorted by Aria cytometer. The knockdown efficiency was verified by real-time PCR. shRNA sequences were as follows: Control-shRNA, 5’-CCGGcaacaagatgaagagcaccaaCTCGAGttggtgctcttcatcttgttgTTTTTG-3’ and 5’-AATTCAAAAAcaacaagatgaagagcaccaaCTCGAGttggtgctcttcatcttgttg-3’; RyR1-shRNA, 5’-CCGGcgtcgcatagaacggatctatCTCGAGatagatccgttctatgcgacgTTTTTG-3’and 5’-AATTCAAAAAcgtcgcatagaacggatctatCTCGAGatagatccgttctatgcgacg-3’; RyR2-shRNA, 5’-CCGGccgctaatgaagccatataaaCTCGAGtttatatggcttcattagcggTTTTTG-3’ and 5’-AATTCAAAAAccgctaatgaagccatataaaCTCGAGtttatatggcttcattagcgg-3’; RyR3-shRNA, 5’-CCGGccgacatggttcagagagaaaCTCGAGtttctctctgaaccatgtcggTTTTTG-3’ and 5’-AATTCAAAAAccgacatggttcagagagaaaCTCGAGtttctctctgaaccatgtcgg-3’; Ahnak-shRNA, 5’-CCGGtgccaccatctactttgacaaCTCGAGttgtcaaagtagatggtggcaTTTTTG -3’ and 5’-AATTCAAAAAtgccaccatctactttgacaaCTCGAGttgtcaaagtagatggtggca-3’; Itpr1--shRNA, 5’-CCGGgcagtaggtaagaagttattaCTCGAGtaataacttcttacctactgcTTTTTG-3’ and 5’-AATTCAAAAAgcagtaggtaagaagttattaCTCGAGtaataacttcttacctactgc-3’; Stim1-shRNA, 5’-CCGGgcagtactacaacatcaagaaCTCGAGttcttgatgttgtagtactgctTTTTTG-3’ and 5’-AATTCAAAAAgcagtactacaacatcaagaaCTCGAGttcttgatgttgtagtactgct-3’; Trpm1-shRNA, 5’-CCGGcggagtgaacatgcagcatttCTCGAGaaatgctgcatgttcactccgTTTTTG-3’ and 5’-AATTCAAAAAcggagtgaacatgcagcatttCTCGAGaaatgctgcatgttcactccg-3’; Trpm4--shRNA, 5’-CCGGgcacatcttcacggtgaacaaCTCGAGttgttcaccgtgaagatgtggTTTTTG-3’ and 5’-AATTCAAAAAgcacatcttcacggtgaacaaCTCGAGttgttcaccgtgaagatgtgc-3’; Trpv2--shRNA, 5’-CCGGccaaggaacttgtttctatttCTCGAGaaatagaaacaagttccttggTTTTTG-3’ and 5’-AATTCAAAAAccaaggaacttgtttctatttCTCGAGaaatagaaacaagttccttgg-3’; Cacna2d4--shRNA, 5’-CCGGtaggaacgcaatggatattaaCTCGAGttaatatccattgcgttcctaTTTTTG-3’ and 5’-AATTCAAAAAtaggaacgcaatggatattaaCTCGAGttaatatccattgcgttccta-3’.

For FoxP3 overexpression, pLVX-IRES-mcherry vectors were a gift from Dr. Xiaohua Shen of School of Medicine, Tsinghua University. For vector construction, FoxP3 was amplified by PCR using cDNA from Treg total RNA as template with the following primers: forward primer 5’-ATCGCTCGAGATGCCCAACCCTAGGCCA-3’ and reverse primer5’-ATCGGAATTCTCAAGGGCAGGGATTGGA-3’. The amplified fragment was gel purified, digested (XhoI and EcoRI) and cloned into the pLVX-IRES-mcherry plasmid, resulting in pLVX-FoxP3-IRES-mcherry construct. Lentivirus production harboring pLVX-FoxP3-IRES-mcherry and FoxP3-overexpressed cell line generation was performed according to above knockdown protocol. FoxP3 expression was verified by Real Time-qPCR.

To overexpress RyR2, we attempted CRISPRa system. All associated vectors including pLenti-EF1a-dCas9-VP64-blast, pLenti-sgRNA(MS2)-EF1a-zeo and pLenti-MS2-P65-HSF1-2A-Hygro were a gift from Dr. Qiaoran Xi of school of life science, Tsinghua University. BleoR sequence in pLenti-sgRNA(MS2)-EF1a-zeo plasmid was replaced with EGFP to generate Lenti-sgRNA (MS2)-EF1a-EGFP backbone plasmids. For designing and cloning sgRNA, in brief, 20-nucleotide gRNA sequences targeting the promoter of RyR2 were designed with the CRISPR design prediction tool (http://crispr.mit.edu/). Selected gRNAs were cloned into Lenti-sgRNA (MS2)-EF1a-EGFP backbone plasmid by following the SAM target sgRNA cloning protocol. We generated sgRNA insertion by annealing each oligos: sgRNA-2, 5’-CACCGCCTCCGGGCCGCCAAACCCG-3’ and 5’-AAACCGGGTTTGGCGGCCCGGAGGC-3’; sgRNA-4, 5’-CACCGGTGCCCTTCCTGACCTCAAG-3’ and 5’-AAACCTTGAGGTCAGGAAGGGCACC-3’; sgRNA-5, 5’-CACCGAGGAGCTCAGCTTCCCGCTG-3’ and 5’-AAACCAGCGGGAAGCTGAGCTCCTC-3’.

The annealing reaction was performed using T4 polynucleotide kinase (NEB) under the following conditions: 37°C for 30 min; 95°C for 5 min; ramp to 25°C at 5°C/min. Diluted annealing product (1:10) was mixed with backbone vector at 1:2.5 mass ratio in Golden Gate reaction using BsmBI restriction endonuclease (NEB) and T4 DNA ligase (NEB). The program was performed on a thermal cycler: 37°C for 5 min, 20°C for 5 min and repeated for 30 cycles; 60°C for 5 min. Through the above steps, gRNA fragment was inserted between the U6 promoter and sgRNA-MS2 scaffold to result in the pLenti-sgRyR2 (MS2)-EF1a-EGFP construct. Then pLenti-EF1a-dCas9-VP64-blast, pLenti-sgRyR2(MS2)-EF1a-EGFP and pLenti-MS2-P65-HSF1-2A-Hygro were co-transfected into MC38 cells and RyR2 expression was verified by Real Time-qPCR.

### Atomic Force Microscopy-Based Single Cell Force Spectroscopy

The experiments were performed as previously described using a JPK CellHesion unit (*2, 39*). In brief, to measure T-DC adhesion forces in bicellular system, DC2.4 cells were cultured on untreated glass disks. T cells were treated with 200 U/ml recombinant human IL-2 overnight. The disks were moved into an AFM-compatible chamber and mounted on to the machine stage. A clean cantilever was coated with CellTak (BD), and then used to glue individual T cells added to the disk. The AFM cantilever carrying a single T cell was lowered to allow T cell contact with an individual DC and to interact for 15 s before being moved upwards, until two cells were separated completely. The force curves were acquired. The process was then repeated. For triple cell system, DC2.4 cells cultured on glass disks were pulsed with 100 μg/ml soluble OVA protein for 4 h before experiments. IL-2-treated Treg/Tconv cells overnight were stained with 10 μM CFSE, and DC2.4 cells were incubated with these fluorescently labeled Treg or Tconv cells for about 20 min before unlabeled OT-II T cells were added. Treg/Tconv cell-DC couples identified with an UV flashlight were then approached by the cantilever tip carrying an OT-II T cell. Treg/Tconv–mediated suppression of OT-II–DC adhesion was assayed. In each cycle, the AFM cantilever carrying a single T cell was lowered by 0.5–2 μm increments until the first force curve was generated. The T cell on the cantilever was then allowed to interact with the DC for 15 s before being moved upwards, until two cells were separated completely. The incubator chamber in which the machine was housed was conditioned at 37°C and at 5% CO_2_. In all experiments, a minimum of 14 force curves were collected for further analysis. The force curves were processed using the JPK image processing software. Only round and robust cells were selected for AFM gluing. For each SCFS experiment, a pair of T-DC was used to generate force readings from each up and down cycle over a period of several minutes; these readings are plotted. At least three such pairs were used for each condition.

### Suppressive Function in vitro

10^4^ purified DCs from splenocytes were pulsed with 2 μg/ml OVA_323-339_ peptide, then suppressor cells (2×10^4^ Treg or RyR2^+/+^ Tconvs or RyR2^-/-^ Tconvs or RyR2 knockdown Tconvs) were added onto DCs to occupy the latter for 30 min. CD25^-^ OTII Tconv cells were stained by CellTrace CFSE (Thermo Fisher Scientific, C34554). 2×10^4^ OTII Tconv cells were mixed in DC-suppressor cell culture to compete OVA-loaded DC occupied by suppressor cells in a 96-well U bottom plate. The proliferation of OTII T cells was assessed by CFSE dilution by Fortessa flow cytometry (BD Biosciences). Inhibition percentage was calculated from (1-proliferation%), then normalized by taking no Treg group as 0% inhibition and Treg group as 100% inhibition.

### RNAseq and ATACSeq

For transcriptome comparison, total RNA of Tconv and Treg cells cultured overnight from wildtype or CKO mice were extracted and sequenced (Annoroad, Beijing, China). Anti-CD3/CD28 activated cells were sequenced as well. For chromatin opening comparison, Tn5-based libraries were built from fixed cells and sequenced (TruePrep DNA Library Kit TD501, Vazyme, Nanjing, China). RNAseq FPKM files were correlated with Pearson correlation method, then hierarchy clustering was calculated and heatmap was plotted with Euclidean Distance. ATACseq data were analyzed with deeptools (version 2.0) then correlated and clustered by Spearman correlation. DEGs were visualized with log10FPKM.

### Visualization and Analysis of ChIP-seq Data

Visible ChIP-seq data were downloaded from GEO Dataset with GEO ID: FoxP3 in Tconv GSM989036, FoxP3 in Tconv cells transduced to express flag-FoxP3 GSM989034. Visualization of ChIP-seq data were facilitated by IGV (v2.4.14) with mouse reference genome mm8. Gene tracks were generated as screen shot in specific gene loci.

### Dual-Luciferase Report Assay

Dual-luciferase report assay was established as described previously (*40*). Murine RyR2 reporter plasmid was constructed by subcloning 1500 bp of the RyR2 promoter into a luciferase expression pGL3 vector. The whole promoter sequence and truncated ones were synthesized and the plasmids were confirmed by sequencing. 1.25×10^5^ FoxP3-overexpressed-3T3, Renca or A20 cells were co-transfected with 300 ng RyR2 reporter and Renilla luciferase reporter plasmids using Neofect (Neofect, TF201201). 36 hrs after transfection, cell lysates were prepared and analyzed using the Dual-Luciferase Report Assay System (Promega).

### Intra-vital Imaging

For basal contact dynamics between T cells and dendritic cells in vivo, RyR2^+/+^ Tconv (Celltrace Far Red labelled), RyR2^-/-^ Tconv (TAMRA labelled) and WT Treg (Far Red-TAMRA duo-labelled) cells were i.v. transferred into CD11c-DTR-eGFP transgenic mice, at 1:1:1 ratio, 3×10^6^ cells each. 12∼18 hours after transfer, inguinal lymph nodes were exposed and imaged at 37°C environment in intrafollicular zone (IFZ). 4D (3D stack + time) videos were imported into Bitplane Imaris software and analyzed. DC channel (eGFP) was calculated as surfaces then set as reference plane for T cells migration. Contacts were designated as T cell-DC distance <1 μm, and contact duration was designated as continuous contact until release (T cell-DC distance >1 μm). For antigen-specific interactions, 6 hrs after Far Red OT-II Tconv cells transfer, 50 μg OVA_323-339_ + 1 μg LPS was injected subcutaneously in right abdomen. Both draining inguinal lymph node (right) and control node (left) were imaged at 12∼18 hrs. For contact disruption of OT-II and DCs by suppressor cell occupation (RyR2^+/+^ Tconv, RyR2^-/-^ Tconv, WT Treg, OT-II Treg and RyR2^-/-^ OT-II Tconv), suppressor cells were labelled with 5 μM CellTrace Yellow (Thermo, USA), and OT-II responder cells were labelled with 5 μM CellTrace FarRed (Thermo, USA). 1:1 mixture of both cells (5×10^6^ each) was i.v. transferred into CD11c-DTR-eGFP transgenic mice. For calcium fluctuation experiments, OT-II responder cells were first labelled with FarRed then loaded with 5 μM calcium indicator FuraRed and transferred. Videos were analyzed with Imaris (Bitplane, Oxford) and contacts between DC and responder T cells with or without suppressor cell occupation were analyzed.

### Mixed Chimera

CD45.1 mice were bred with CD45.2 C57BL/6 mice to generate CD45.1/2 mice expressing both the allelic variants. Mixed bone marrow chimeras were generated by intravenously injecting 5×10^5^ CD45.2 CKO bone marrow cells in conjunction with an equal number of CD45.1 control bone marrow cells into lethally X-ray irradiated (two doses of 5.5 Gy, two hours apart) 5-week CD45.1/2 recipient mice. Mice were sacrificed after 8 weeks after bone marrow transplantation. Thymus, spleen and lymph node from mixed bone marrow chimeras were harvested for characterization of lymphocyte development.

### TCR repertoire

To assess whether RyR2 deletion affects T cell thymic selection, T cell receptor Vβ chains of splenic CD4^+^ T cells from CKO and control mice were assessed using a commercial mouse TCR Vβ screening panel (BD Biosciences). Flow cytometry data collection was performed on an LSR II (BD Biosciences). Data on the expression of TCR vβ chains in splenic CD4^+^ T cells for peripheral TCR usage and the frequencies of CD4^+^ Tconvs and Tregs expressing commonly utilized TCR Vβ chains were analyzed using FlowJo V10.

### H&E Histology

Mice were sacrificed by necropsy. Skin, ear, liver, and other tissues were all fixed in 4% neural buffered formalin at 4°C over 48 h before processing. Then samples were embedded in paraffin. Five to six micrometer thick slides were cut. All slides were stained with hematoxylin and eosin. Inflammation score: 0, no inflammatory infiltration; 1, sparse infiltration; 2, obvious infiltration; 3, massive infiltration with normal tissue morphology; 4, massive infiltration with tissue deform; 5, complete tissue destruction.

### HSV-1 Infection Model

Herpes infection was induced by 10^6^ pfu HSV-1 (F strain) in 20 μL PBS into hinder footpads at 0. day At 3 dpi, cells (Treg cells, RyR2^+/+^ Tconv cells or RyR2^-/-^ Tconv cells) were adoptively transferred into footpad with 2×10^5^ cells/20 μL PBS. Virus titer was tested on 7 dpi with homogenized footpad tissue on Vero cells. For DTH, right footpad was re-challenged with UV-inactivated HSV-1 (10^6^ pfu/20 μL PBS) at 6 dpi, then footpad swelling was measured at 7 dpi with left footpad as control.

### Asthma Model

Airway inflammation was induced by ovalbumin (OVA). At day 0 and day 14, two *i*.*p*. injections of OVA plus alum adjuvant (100 μg+4 mg in 200 μL PBS) were given to sensitize mice. Intra-tracheal OVA re-challenge (50 μg in 50 μL PBS) were repeatedly given at back of the tongue on days 21, 23 and 25. At day 23, 10^6^ cells (Treg cells, RyR2^+/+^ Tconv cells or RyR2^-/-^ Tconv cells) were adoptively transferred through *i*.*v*. in 100 μL PBS. BALF infiltrates and histology were analyzed at day 32.

### DSS-Induced Colitis Model

DSS-induced murine experimental colitis was established as described previously (*41, 42*). Briefly, 3×10^6^ RyR2^-/-^ Tconvs, WT Tconvs or Tregs were transferred into 6-week-old male C57BL/6 mice *i*.*v*.. The second day, colitis was induced by oral administration of 4% DSS (w/v) (Yeasen, MW = 36,000-50,000 Da) in drinking water for 7 days followed by normal drinking water. Normal control mice were treated with PBS and were given normal drinking water. The mice were sacrificed on day 10, colons were dissected and colon length was measured. Then the colons were fixed at 4°C over 48 hrs in 4% PFA for subsequent H&E staining. The colon sections were examined by 3DHISTECH Pannoramic SCAN (3DHISTECH).

### MC38 Tumor Model

The grafted tumor mouse model was established as previously described (*43*). Briefly, MC38 colon cancer cells were washed twice in PBS, and 5 × 10^5^ MC38 cells in 200 μL of PBS were coinjected with 10^6^ T cells s.c. into the abdomen of 6-week-old male C57BL/6 mice after barbering. Tumor growth was monitored using a slide caliper every 4∼5 days from day 7 after tumor injection. The volume was calculated as (length×width×width) / 2. Group 1: MC38+PBS; Group 2: MC38+Tconvs; Group 3: MC38+Tregs; Group 4: MC38+RyR2^-/-^ Tconvs;. There were ten mice in each group.

### Rescue of Scurfy Mice

The rescue of FoxP3-deficient mice was established as previously described (*44, 45*). Scurfy mice’s genotypes were analyzed at birth with pamprodactylous-clipped tissue routinely. First adoptive transfer of 5×10^6^ purified cells (Treg cells, FoxP3-deficient Scurfy Tconv cells or CD25^-^RyR2^-/-^ Tconv cells) was performed on day 2 or 3 of life in 50 uL PBS for i.p. injection into newborn syngeneic Scurfy mice. Then, in the first two weeks, cells transfer was performed every three or four days, then once every two weeks. When transferring, the body weight of Scurfy and male WT littermates was recorded and survival was monitored. Genotyping for the sf mutant gene was conducted by PCR and verified by sequencing. Primers for FoxP3 PCR were 5’-CATCCCACTGTGACGAGATG-3’ and 5’-ACTTGGAGCACAGGGGTCT-3’. For the histology, PBS and FoxP3^-^ Tconv-infused mice were analyzed on week 3, and those from Treg and RyR2^-/-^ Tconv-infused were on week 8-12. Skin, ear, liver, and other tissues were all fixed for H&E staining.

### Statistial analysis

Numbers of experimental repeats are shown in the figure legend. Student’s t test was used for comparing endpoint means of different groups. Data were always presented as mean ± SEM, unless indicated otherwise. Calculation and graphing were done with Prism (GraphPad). P value < 0.05 was considered significant. *, P < 0.05; **, < 0.01; ***, < 0.001; ****, < 0.0001; N.S., not significant.

## Supplementary Materials

Figs. S1 to S6

## Acknowledgments

We thank Drs. Kenneth Rock, Xu Tan, Guangyu An, and Wei Guo for providing cell lines, Drs. Qiaoran Xi and Xiaohua Shen for providing plasmids, and Dr. Bin Wang for providing HSV-1. We thank the Microscopy and Imaging Facility and Live Cell Imaging Facility of the University of Calgary for technical assistance. We also thank Imaging Core Facility, Technology Center for Protein Sciences of Tsinghua University for assistance. We thank Mr. Guohua Yuan for critical reading.

## Author Contributions

X.W (Xiaobo Wang) and J.M performed all the experiments unless noted otherwise. X.W (Xiaobo Wang) and S.G. designed and performed all animal experiments with help from J.G. and Z.Z.. J.M and N.K performed AFM-related experiments. X.L. performed ChIP-qPCR and in vitro suppression. X.X. performed calcium imaging. H.L and Y.X. (Ying Xu) performed shRNA screening of knockdown cells. B.Z. analyzed and visualized ChIP-seq data. Y.X. (Yanni Xu) and X.W. (Xin Wang) performed RyR2 protein detection. X.S. and S.G. performed intra-vital imaging assay. D.Z. and X.W. (Xiaoting Wang) performed cytokine detection. X.M. and N.N. performed transcription analysis of Calcium channel genes. R.S. performed histology analysis. R.W. assisted RyR2 function analysis. J.G. provided Scurfy and Treg biology expertise. W.C. provided expert insight in RyR2 regulation. X.Z. provided critical review on Treg biology. T.X. provided biophysical mechanism on cell adhesion. H.Q. provided overall critical insight. X.H. provided expert insight in gene regulation of RyR2. Y.S. was responsible for the overall design and wrote the manuscript with inputs from X.H, H.Q, and R.W.

