## Supplemental figure S1-S6 for "FoxP3-mediated blockage of ryanodine receptor 2 is the molecular basis for the contact-based suppression by regulatory T cells"

### Fig. S1

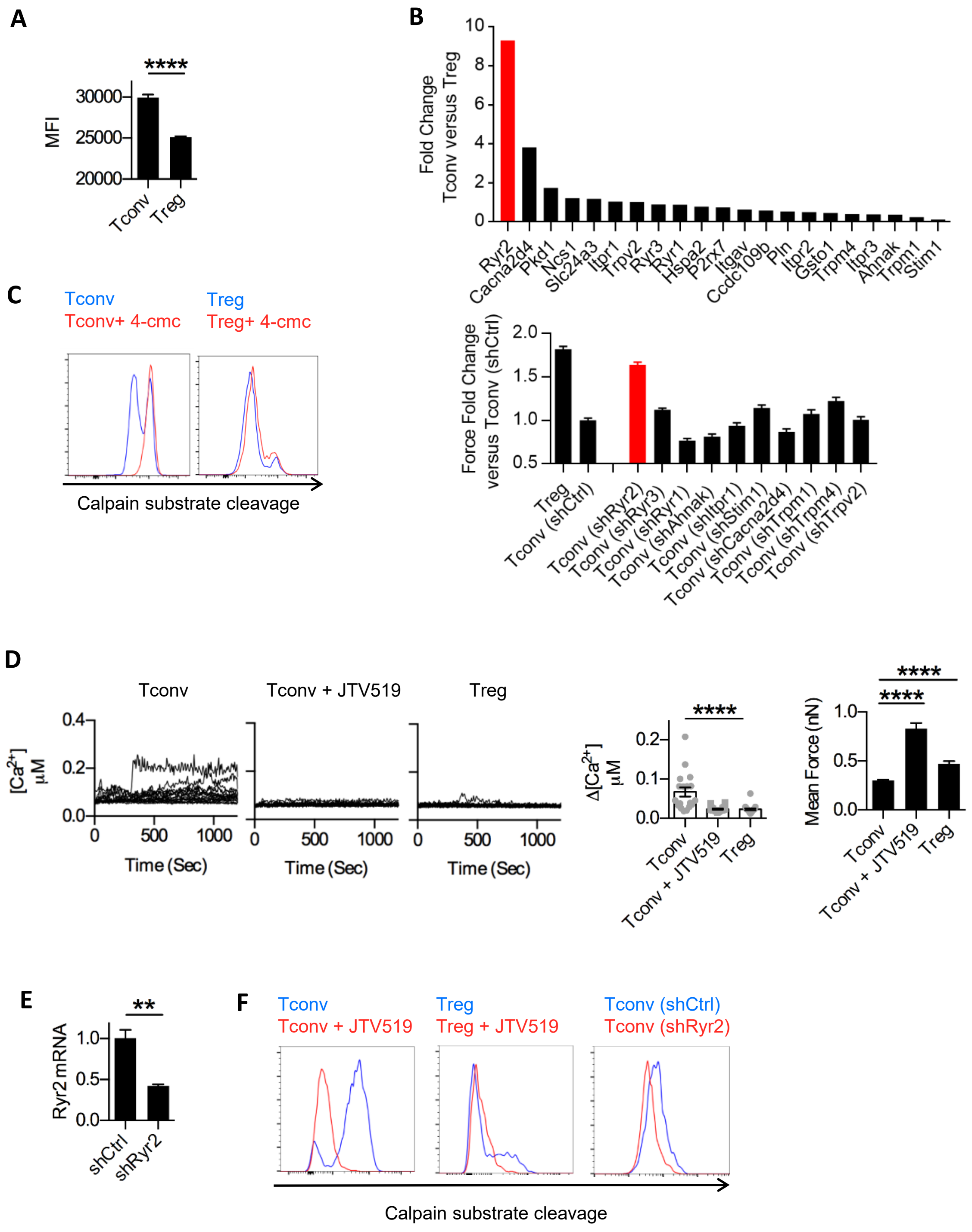

A
****
MFI
30000
25000
20000
Tconv
Treg
B
Fold Change
Tconv versus Treg
10
8
6
4
2
0
Ryr2
Cacna2d4
Pkd1
Ncs1
Slc24a3
Itpr1
Trpv2
Ryr3
Ryr1
Hspa2
P2rx7
Itgav
Ccdc109b
Pln
Itpr2
Gsto1
Trpm4
Itpr3
Ahnak
Trpm1
Stim1
C
Tconv
Tconv+ 4-cmc
Treg
Treg+ 4-cmc
Calpain substrate cleavage
Force Fold Change
versus Tconv (shCtrl)
2.0
1.5
1.0
0.5
Treg
Tconv (shCtrl)
Tconv (shRyr2)
Tconv (shRyr3)
Tconv (shRyr1)
Tconv (shAhnak)
Tconv (shItpr1)
Tconv (shStim1)
Tconv (shCacna2d4)
Tconv (shTrpm1)
Tconv (shTrpm4)
Tconv (shTrpv2)
D
Tconv
Tconv + JTV519
Treg
[Ca2+] μM
0.4
0.2
0.0
0
500
1000
Time (Sec)
Δ[Ca2+] μM
****
0.2
0.1
0.0
Tconv
Tconv + JTV519
Treg
Mean Force (nN)
****
****
1.0
0.5
0.0
E
**
Ryr2 mRNA
1.0
0.5
0.0
shCtrl
shRyr2
F
Tconv
Tconv + JTV519
Treg
Treg + JTV519
Tconv (shCtrl)
Tconv (shRyr2)
Calpain substrate cleavage

**Fig. S1. Tregs lack of RyR2-dependent basal calcium oscillations.** (**A**) Quantification of the mean fluorescence intensity (MFI) of $Ca^{2+}$ signals of Tconvs and Tregs under resting state with Fluo 4. N=5. (**B**) Upper panel, qPCR analysis of $Ca^{2+}$ regulation-associated protein levels between Tregs and Tconvs, ranked by the fold of differences. Lower panel, relative fold change of mean forces. IL-2-treated Treg cells and shRNA samples of Ryr2, Ryr1, Ryr3, Ahnak, Itpr1, Stim1, Cacna2d4, Trpm1, Trpm4 and Trpv2 genes were normalized to control Tconv cells (shCtrl). Force were measured with DC2.4 cells. N=3. (**C**) Analysis of calpain activities in Tconv (left) and Treg (right) cells treated with 4-CMC as determined by CMAC digestion. N=4. (**D**) Resting Tconv loaded with Fluo-4 AM were treated with 5 μM JTV519 for 30 min then Fluo-4 fluorescence signal over time was recorded. Untreated Tconvs and Tregs were as control. The change of intracellular free $Ca^{2+}$ concentration [$Ca^{2+}$] over time were shown. Corresponding amplitude was shown in the middle. n=20, N =5. Adhesion force of Tconv treated with JTV519 was shown in the right. N=3. (**E**) Analysis of Ryr2 gene shRNA knockdown efficiency in Tconv cells isolated from FoxP3-GFP mice using qPCR. N=3. (**F**) Analysis of the calpain activities in JTV519-treated Tconvs (left), Tregs (middle) and RyR2 knockdown Tconvs (right) cells by CMAC digestion. N=4. Here, **, < 0.01; ****, < 0.0001.

Fig. S2

A

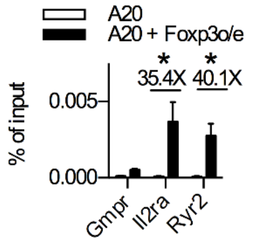


**Fig. S2. RyR2 is transcriptionally silenced by FoxP3.** (**A**) ChIP-qPCR analysis was performed in FoxP3-overexpressed A20 cells to examine FoxP3 enriched binding of Ryr2 promotor region, with Gmpr as negative control and IL2ra as positive control. N=3. Here, *, < 0.05.

Fig. S3

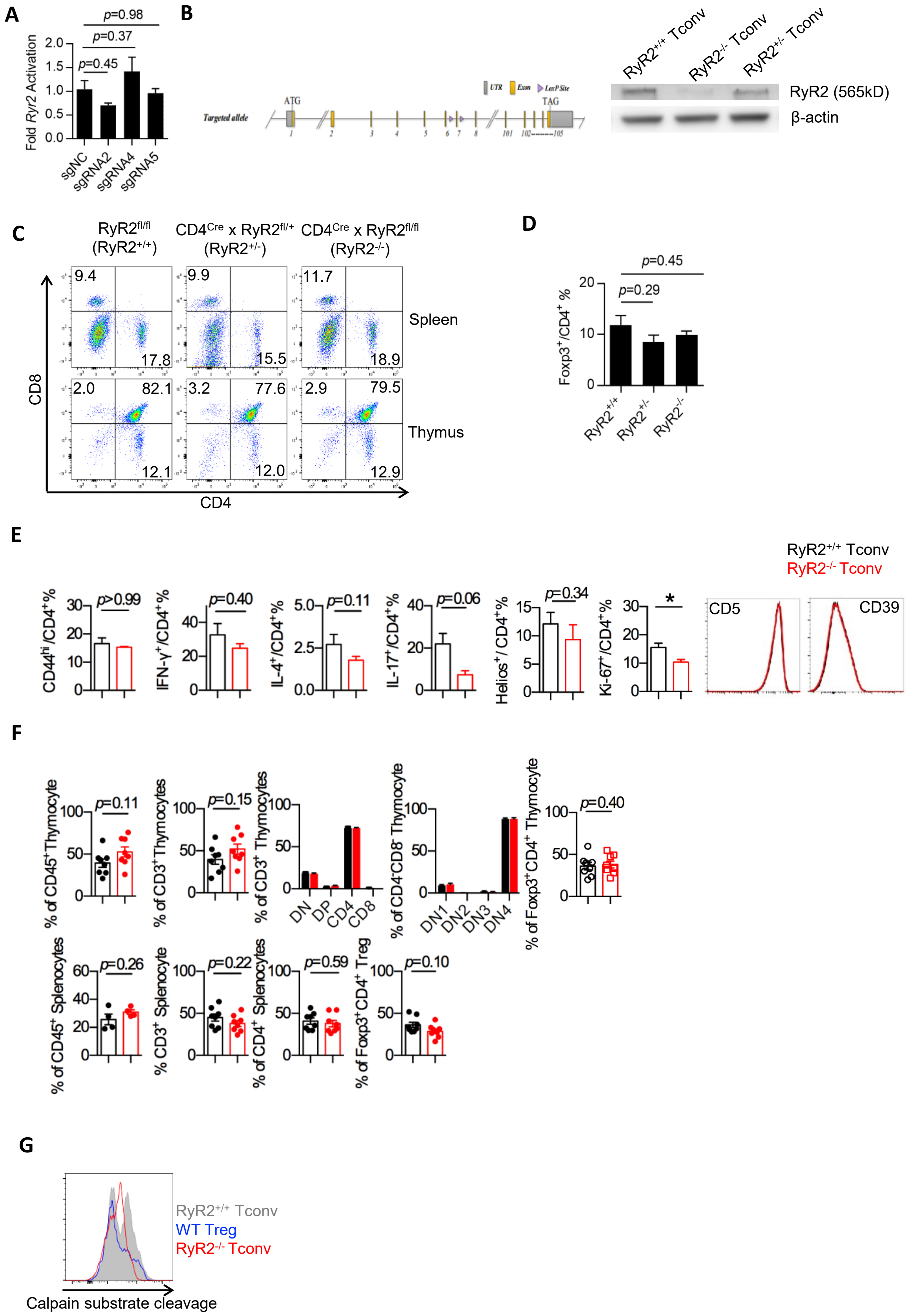

A
p=0.98
p=0.37
p=0.45
Fold Ryr2 Activation
2.0
1.5
1.0
0.5
0.0
sgNC
sgRNA2
sgRNA4
sgRNA5
B
Targeted allele
ATG
TAG
UTR
Exon
LoxP Site
RyR2+/+ Tconv
RyR2-/- Tconv
RyR2+/- Tconv
RyR2 (565kD)
β-actin
C
RyR2fl/fl
(RyR2+/+)
CD4Cre x RyR2fl/+
(RyR2+/-)
CD4Cre x RyR2fl/fl
(RyR2-/-)
9.4
17.8
9.9
15.5
11.7
18.9
Spleen
2.0
82.1
12.1
3.2
77.6
12.0
2.9
79.5
12.9
Thymus
CD8
CD4
D
p=0.45
p=0.29
Foxp3+/CD4+ %
20
10
0
RyR2+/+
RyR2+/-
RyR2-/-
E
RyR2+/+ Tconv
RyR2-/- Tconv
CD44hi/CD4+%
p>0.99
IFN-γ+/CD4+%
p=0.40
IL-4+/CD4+%
p=0.11
IL-17+/CD4+%
p=0.06
Helios+/CD4+%
p=0.34
Ki-67+/CD4+%
*
CD5
CD39
F
% of CD45+Thymocyte
p=0.11
% of CD3+Thymocytes
p=0.15
% of CD3+ Thymocytes
DN
DP
CD4
CD8
% of CD4-CD8- Thymocyte
DN1
DN2
DN3
DN4
% of Foxp3+CD4+ Thymocyte
p=0.40
% of CD45+ Splenocytes
p=0.26
% CD3+ Splenocyte
p=0.22
% of CD4+ Splenocytes
p=0.59
% of Foxp3+CD4+ Treg
p=0.10
G
RyR2+/+ Tconv
WT Treg
RyR2-/- Tconv
Calpain substrate cleavage

**Fig. S3. RyR2-deficiency minimally affects T cell development.** (**A**) Fold activation of Ryr2 transcripts in MC38 cells by three sgRNAs using an engineered CRISPR-Cas9 complex (CRISPRa system) were shown. qPCR analysis was performed. N=3. (**B**) The construction of conditional knockout mice with intended Cre-dependent deletion of exon 7 in Ryr2 gene (left) was shown. RyR2 of $CD4^+$ T cells was deleted in CKO mice examined by Western Blot (right). (**C**) Distribution of $CD4^+$ and $CD8^+$ T cells in spleen and thymus of $Ryr2^{fl/fl}$ ($RyR2^{+/+}$) mice, CD4-Cre/$RyR2^{fl/+}$ ($RyR2^{+/-}$) mice and CD4-Cre/$Ryr2^{-/-}$ ($RyR2^{-/-}$) mice was detected by flow cytometry. N=3. (**D**) The proportion of $CD4^+FoxP3^+$ cells in $CD4^+$ splenocytes in mice shown in C. n=3, N=3. (**E**) The frequency of $CD44^{hi}$, $IFN\gamma^+$, $IL\text{-}4^+$, $IL\text{-}17^+$, $Helios^+$ and $Ki67^+$ T cells and CD39/CD5 expression levels of $CD4^+$ splenic T cells from CKO mice were measured by FACS, N=3. (**F**) Mixed bone marrow chimera. Thymus and spleens from mixed chimeras were harvested for characterization. Shown are $RyR2^{+/+}$ CD45.1 and $RyR2^{-/-}$ CD45.2 percentages in $CD45^+$, $CD3^+$, $CD4^-CD8^-$, $CD4^+CD8^+$, $CD4^+$, $CD8^+$ thymocytes, and $CD25^-CD44^+$ (DN1), $CD25^+CD44^+$ (DN2), $CD25^+CD44^-$ (DN3), $CD25^-CD44^-$ (DN4) $CD4^-CD8^-$ thymocytes, $FoxP3^+$ $CD4^+$ thymocytes (upper), $CD45^+$, $CD3^+$, $CD4^+$, $CD8^+$ and $FoxP3^+$ $CD4^+$ splenocytes (lower). One dot represents one mouse. N=2. (**G**) Calpain activities in Treg, $RyR2^{+/+}$ and $RyR2^{-/-}$ Tconv cells were measured by using CMAC digestion. N=3. Here, *, $< 0.05$.

### Fig. S4

A

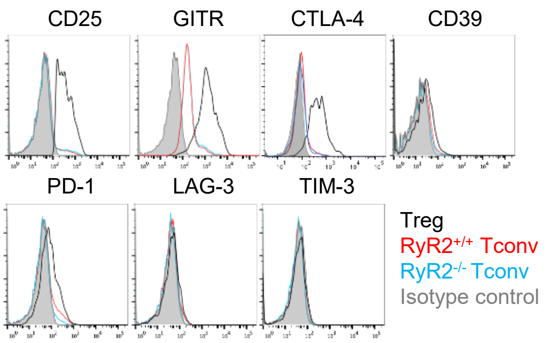


B

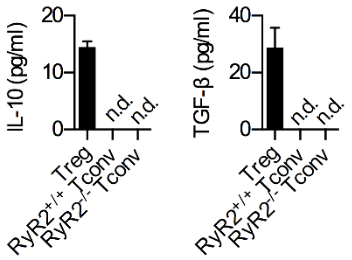


C

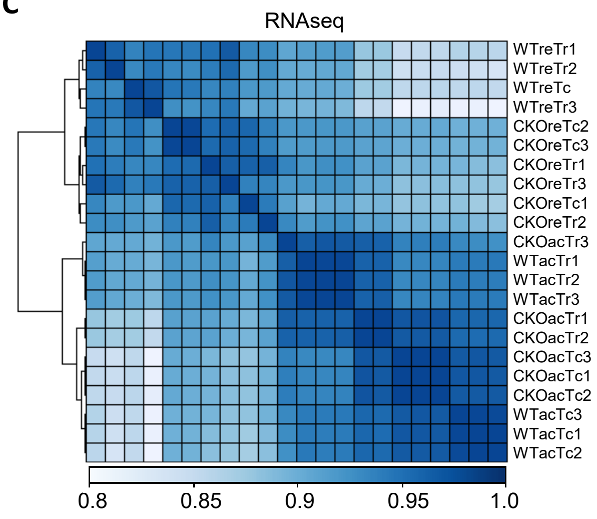


D

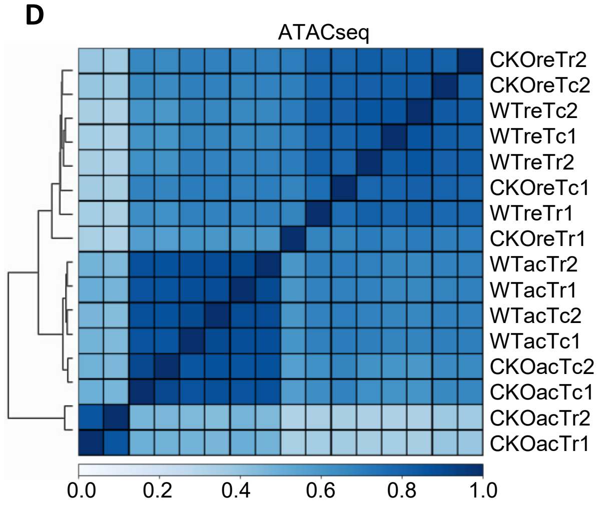


E

| GeneName | WTreTc | CKOreTc | WTreTr | WTacTc | CKOacTc | WTacTr |
|---|---|---|---|---|---|---|
| Foxa1 | 0.00 | 0.04 | 0.33 | 0.00 | 0.00 | 0.02 |
| Foxa2 | 0.00 | 0.00 | 0.03 | 0.00 | 0.00 | 0.00 |
| Foxa3 | 0.00 | 0.00 | 0.04 | 0.04 | 0.16 | 0.09 |
| Foxb1 | 0.00 | 0.00 | 0.00 | 0.51 | 0.02 | 0.14 |
| Foxc1 | 0.00 | 0.01 | 0.04 | 0.05 | 0.05 | 0.13 |
| Foxc2 | 0.11 | 0.10 | 0.06 | 0.00 | 0.00 | 0.00 |
| Foxd1 | 0.00 | 0.00 | 0.00 | 0.00 | 0.00 | 0.00 |
| Foxd2 | 0.72 | 0.49 | 0.96 | 0.50 | 0.25 | 0.62 |
| Foxd2os | 1.87 | 0.81 | 1.25 | 1.08 | 0.78 | 1.78 |
| Foxd3 | 0.00 | 0.00 | 0.00 | 0.00 | 0.03 | 0.00 |
| Foxd4 | 0.00 | 0.00 | 0.00 | 0.00 | 0.00 | 0.00 |
| Foxe1 | 0.00 | 0.00 | 0.00 | 0.05 | 0.00 | 0.00 |
| Foxf1 | 0.02 | 0.03 | 0.03 | 0.00 | 0.00 | 0.02 |
| Foxf2 | 0.03 | 0.00 | 0.00 | 0.15 | 0.03 | 0.08 |
| Foxh1 | 0.00 | 0.00 | 0.18 | 0.00 | 0.00 | 0.18 |
| Foxi1 | 0.00 | 0.00 | 0.03 | 0.03 | 0.03 | 0.00 |
| Foxi3 | 0.00 | 0.00 | 0.00 | 0.00 | 0.00 | 0.00 |
| Foxj1 | 0.05 | 0.23 | 0.35 | 2.54 | 0.78 | 1.28 |
| Foxj2 | 16.36 | 12.99 | 15.75 | 6.73 | 6.08 | 9.45 |
| Foxj3 | 8.29 | 10.99 | 12.42 | 8.62 | 8.99 | 9.62 |
| Foxk1 | 31.83 | 39.18 | 22.07 | 15.92 | 16.72 | 15.77 |
| Foxk2 | 12.98 | 11.55 | 11.92 | 13.83 | 14.25 | 16.24 |
| Foxl2 | 0.00 | 0.04 | 0.00 | 0.00 | 0.00 | 0.00 |
| Foxl2os | 0.00 | 0.02 | 0.00 | 0.04 | 0.00 | 0.05 |
| Foxm1 | 2.41 | 2.21 | 2.50 | 2.23 | 1.33 | 4.16 |
| Foxn1 | 0.00 | 0.00 | 0.00 | 0.00 | 0.00 | 0.02 |
| Foxn2 | 18.86 | 15.86 | 15.91 | 16.00 | 16.13 | 11.96 |
| Foxn3 | 17.46 | 18.74 | 16.34 | 11.62 | 11.97 | 13.44 |
| Foxo1 | 25.96 | 32.56 | 29.51 | 16.58 | 14.61 | 17.43 |
| Foxo3 | 15.16 | 18.25 | 16.18 | 10.69 | 11.33 | 13.23 |
| Foxo4 | 17.51 | 27.64 | 25.38 | 6.23 | 5.38 | 10.87 |
| Foxo6 | 0.00 | 0.05 | 0.00 | 0.00 | 0.00 | 0.00 |
| Foxp1 | 20.14 | 29.17 | 18.10 | 22.64 | 24.43 | 19.06 |
| Foxp2 | 0.01 | 0.00 | 0.00 | 0.01 | 0.00 | 0.02 |
| Foxp3 | 0.62 | 2.83 | 118.29 | 1.07 | 1.96 | 123.45 |
| Foxp4 | 22.00 | 22.78 | 27.00 | 24.49 | 27.40 | 29.55 |
| Foxq1 | 0.04 | 0.30 | 0.66 | 0.15 | 0.04 | 0.01 |
| Foxr1 | 0.00 | 0.00 | 0.00 | 0.00 | 0.00 | 0.00 |
| Foxred1 | 7.08 | 5.98 | 6.41 | 7.03 | 8.89 | 6.83 |
| Foxred2 | 2.89 | 2.28 | 2.24 | 5.21 | 4.84 | 6.05 |
| Foxs1 | 0.14 | 0.00 | 0.00 | 0.17 | 0.00 | 0.00 |

F

| GeneName | WTreTc | CKOreTc | WTreTr | WTacTc | CKOacTc | WTacTr |
|---|---|---|---|---|---|---|
| Ryr1 | 0.13 | 0.05 | 0.04 | 0.13 | 0.10 | 0.08 |
| Ryr3 | 0.25 | 0.31 | 0.71 | 0.04 | 0.05 | 0.12 |
| Cacna2d4 | 3.59 | 2.26 | 3.80 | 0.40 | 0.46 | 1.32 |
| Ahnak | 40.30 | 22.49 | 59.47 | 9.65 | 8.48 | 25.64 |
| Itpr1 | 6.03 | 6.33 | 5.08 | 4.24 | 3.91 | 4.46 |
| Itpr2 | 23.20 | 19.14 | 24.23 | 7.32 | 5.26 | 11.56 |
| Itpr3 | 37.00 | 37.82 | 29.29 | 6.29 | 5.77 | 11.78 |
| Stim1 | 13.87 | 18.14 | 11.79 | 5.93 | 6.19 | 7.33 |
| Stim2 | 12.84 | 15.45 | 18.11 | 17.89 | 19.05 | 13.99 |
| Trpm1 | 0.13 | 0.10 | 0.06 | 0.00 | 0.00 | 0.16 |
| Trpm2 | 0.20 | 1.29 | 0.13 | 0.00 | 0.00 | 0.01 |
| Trpm3 | 0.00 | 0.00 | 0.00 | 0.00 | 0.00 | 0.00 |
| Trpm4 | 1.46 | 1.34 | 2.10 | 1.09 | 0.84 | 1.80 |
| Trpv1 | 0.00 | 0.00 | 0.00 | 0.00 | 0.00 | 0.00 |
| Trpv2 | 21.08 | 16.50 | 24.14 | 11.51 | 9.89 | 22.63 |
| Trpv3 | 0.04 | 0.00 | 0.00 | 0.00 | 0.00 | 0.00 |
| Trpv4 | 0.06 | 0.12 | 0.02 | 0.05 | 0.02 | 0.13 |

**Fig. S4. RyR2 deficiency per se mediates contact-dependent suppression.** (**A**) Analysis of surface markers potentially associated with Treg functions in RyR2$^{+/+}$ and RyR2$^{-/-}$ Tconvs by flow cytometry. (**B**) IL-10 and TGF-β from Treg, RyR2$^{+/+}$ and RyR2$^{-/-}$ Tconv cells after 72 hrs anti-CD3 plus anti-CD28 stimulation were tested by ELISA. (**C**) Spearman correlation and hierarchy clustering of the RNAseq data. WT or CKO Tconvs and Tregs were sequenced in resting (eg. WTreTc) or anti-CD3/CD28 activated (eg. WTacTc) status. Euclidean distance among RNAseq samples were shown. (**D**) Spearman correlation among ATACseq samples. (**E**) Transcription level of Forkhead family members. FPKM of each gene was shown. (**F**) Transcription level of T cell associated calcium regulators. FPKM of each gene was shown.

Fig. S5

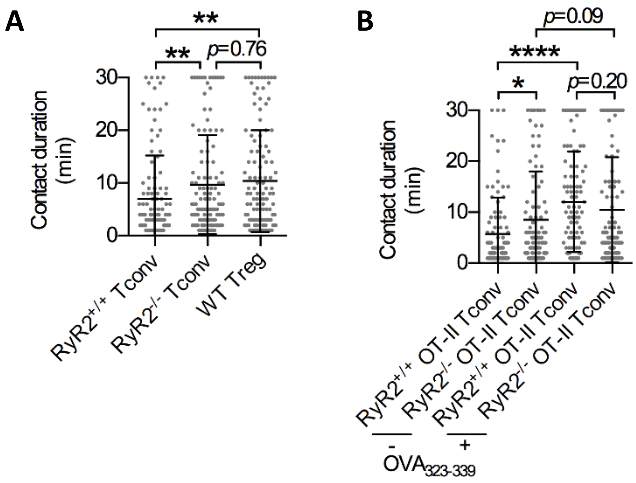


**Fig. S5. RyR2 deficiency-mediated suppression operates in the absence of specific antigen.** (**A**) Basal contact duration of $RyR2^{+/+}$ Tconv, $RyR2^{-/-}$ Tconv and Treg cells to DC in vivo. CD11c-DTR-eGFP transgenic mice were i.v. transferred with labeled $RyR2^{+/+}$ Tconv, $RyR2^{-/-}$ Tconv and WT Treg cells at 1:1:1 ratio. 120 contacts from 30~50 Tconv/Treg cells were analyzed. (**B**) Antigen-specific contact duration of $RyR2^{+/+}$ OT-II Tconv, $RyR2^{-/-}$ OT-II Tconv. CD11c-DTR-eGFP transgenic mice were i.v. transferred with labeled $RyR2^{+/+}$ OT-II Tconv or $RyR2^{-/-}$ OT-II Tconv, then inoculated with $OVA_{323\text{-}339}$ mixed with LPS at right abdomen region. Both draining inguinal lymph node ($OVA_{323\text{-}339}$+) and control node ($OVA_{323\text{-}339}$ free) were analyzed. 100 contacts from 20~50 cells are analyzed. Data are pooled from 3 independent experiments. N=3. Here, *, < 0.05; **, < 0.01; ****, < 0.0001.

Fig. S6

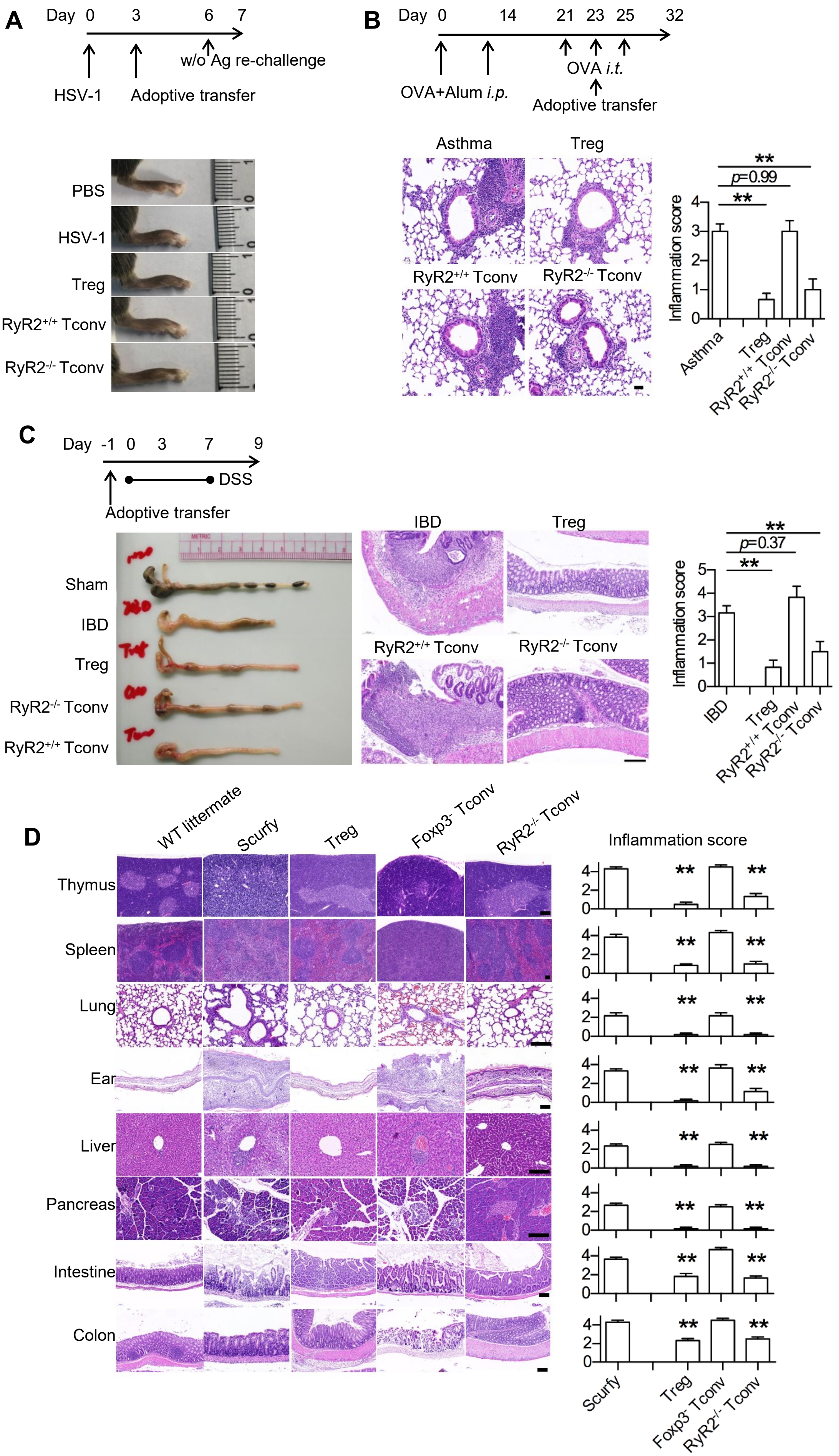

A
Day 0 3 6 7
HSV-1
Adoptive transfer
w/o Ag re-challenge
PBS
HSV-1
Treg
RyR2+/+ Tconv
RyR2-/- Tconv
B
Day 0 14 21 23 25 32
OVA+Alum i.p.
OVA i.t.
Adoptive transfer
Asthma
Treg
RyR2+/+ Tconv
RyR2-/- Tconv
Inflammation score
p=0.99
C
Day -1 0 3 7 9
DSS
Adoptive transfer
Sham
IBD
Treg
RyR2-/- Tconv
RyR2+/+ Tconv
Inflammation score
p=0.37
D
WT littermate
Scurfy
Treg
Foxp3- Tconv
RyR2-/- Tconv
Inflammation score
Thymus
Spleen
Lung
Ear
Liver
Pancreas
Intestine
Colon

**Fig. S6. RyR2-deficient Tconvs are indistinguishable from Tregs in disease models and scurfy rescue.** (**A**) Experimental scheme (upper) and representative footpad swelling images in DTH assay (lower left). (**B**) Sensitization scheme (upper) and representative lung H&E histological sections (lower). Scale bars represent 200 μm. Tissue inflammations were scored as methods stated. N=3. (**C**) Induction scheme of DSS-induced colitis model (left), representative colon images (lower left) and photomicrograph of colon sections (lower right). Scale bars represent 200 μm. (**D**) Representative H&E staining of samples taken from indicated organs or anatomic sites. For WT and Scurfy and $FoxP3^-$ infused Scurfy, samples were taken on week 3. Rescued Scurfy mice were taken on week 8-12. Here, **, < 0.01; ***, < 0.001.
